# Tools for efficient analysis of neurons in a 3D reference atlas of whole mouse spinal cord

**DOI:** 10.1101/2021.05.06.443008

**Authors:** Felix Fiederling, Luke A. Hammond, David Ng, Carol Mason, Jane Dodd

## Abstract

Spinal neurons are highly heterogeneous in location, transcriptional identity and function. To understand their contributions to sensorimotor circuits, it is essential to map the positions of identified subsets of neurons in relation to others throughout the spinal cord (SC), but we lack tools for whole SC sample preparation, imaging and *in toto* analysis. To overcome this problem, we have (1) designed scaffolds (SpineRacks) that facilitate efficient and ordered cryo-sectioning of the entire SC in a single block, (2) constructed a 3D reference atlas of adult mouse SC and (3) developed software (SpinalJ) to register images of sections and for standardized analysis of cells and projections in atlas space. We have verified mapping accuracies for known neurons and demonstrated the usefulness of this platform to reveal unknown neuronal distributions. Together, these tools provide high-throughput analyses of whole mouse SC and enable direct comparison of 3D spatial information between animals and studies.

## Introduction

The spinal cord (SC) integrates sensorimotor signals that ultimately produce the precise patterns of motor activity that control all body movements. Spinal neurons receive and process somatosensory signals from skin, muscles, joints and viscera and direct local and brain-derived motor commands. A huge body of work has provided a good understanding of the roles of major SC neuron types in sensorimotor processing and there are now many examples of well characterized neurons whose morphology, position, interconnections and functions are known within defined segments of SC. Emerging from this work is the importance of stereotypic positioning of spinal neurons as a basis of circuit specificity, both dorsoventrally (d-v) in the grey matter within laminae and clusters, and rostrocaudally (r-c), along the axis of the SC, to accommodate distinct body regions. Our current knowledge comes from classical *in vivo* electrophysiology and anatomy and from work exploring the genetic bases of SC development and the specification of cell types (Abraira and Ginty, 2013; Brown, 1982; Gatto et al., 2019; Goulding, 2009; Jessell, 2000; Lai et al., 2016; Rexed, 1954; Sherrington, 1906; Stachowski and Dougherty, 2021). These studies have also produced genetic markers for identifying and tracing cell development and for manipulating neuronal function in subsets of mouse spinal neurons. More recently, advances in single cell transcriptional profiling have revealed a further level of cellular heterogeneity and complexity within the SC (Delile et al., 2019; Häring et al., 2018; Rosenberg et al., 2018; Sathyamurthy et al., 2018) that transcends classical subgroupings. This previously unappreciated subpopulation heterogeneity within the cardinal classes of spinal neurons in turn demands finer delineation of neurons and their connections to understand the cellular architecture of functional circuitry. While molecular insights and the precision of modern genetic tools provide the means to access and label increasingly specific subsets of spinal neurons, positioning such data in relation to other neurons and circuits within the framework of the whole SC has been elusive, as we lack tools for 3D analysis of whole mouse SC.

Visualizing cells and connections in a structure that, in mouse, spans 3-4cm is technically challenging. Traditional histological approaches are prohibitively labor intensive: sectioning the SC of adult mouse at 20μm produces ~1900 sections (Watson et al., 2009); manually collecting, staining and imaging these sections while maintaining r-c order is painstaking and the lack of tools for 3D reconstruction and analysis within a standardized reference limits data comparison. An alternate approach is to image neurons within intact SC. Two-photon microscopy has been used to visualize axons in superficial layers of fixed or unfixed whole SC, but myelinated fiber bundles hinder deep imaging (Hilton et al., 2019; Johannssen and Helmchen, 2013). The recent introduction of tissue clearing (reviewed in Tian and Li, 2020; Ueda et al., 2020) in combination with light sheet microscopy has enabled fast, 3D visualization of large intact samples (Cai et al., 2019; Hillman et al., 2019; Pan et al., 2016; Zhao et al., 2020). However, while clearing has been used effectively to visualize and analyze certain genetically labeled spinal tracts and to assess axon regeneration in SC injury models (Ertürk et al., 2012a; Hilton et al., 2019), complete clarity in mature SC remains elusive because of the abundance of myelin that introduces light scattering (Soderblom et al., 2015). Moreover, solvent based clearing methods are often unsuitable for non-genetic labeling, interfering with endogenous protein fluorescence (Ertürk et al., 2012b; Pan et al., 2016; Qi et al., 2019; Renier et al., 2014), and are incompatible with lipophilic tracers and DNA dyes (Tian and Li, 2020). Aqueous- and hydrogel-based methods that counter these issues have been successful in preserving protein-based fluorescence. However, labeling remains limited by antibody compatibility and low penetration rates (Tian and Li, 2020) and, in fact, tissues such as whole adult SC exceed the size that can be cleared or processed for immunohistochemistry using these protocols (Vigouroux et al., 2017). Thus, there remains a pressing need for alternative methods to acquire whole SC 3D image datasets.

Irrespective of the method of acquisition, resources for analyzing whole SC data are scarce and lag far behind the manifold open-source and commercial tools available for whole brain reconstruction, atlas registration and data interpretation, for example see: (Bakker et al., 2015; Botta et al., 2020; Chon et al., 2019; Eastwood et al., 2019; Friedmann et al., 2020; Oh et al., 2014; Puchades et al., 2019; Shiffman et al., 2018; Tappan et al., 2019; Tyson et al., 2020; Wang et al., 2021). Although 3D reconstruction and analysis of human SC MRI data has been reported, the tools offer only low resolution data registration of larger gray and white matter regions (De Leener et al., 2017; Prados et al., 2016). Thus, to advance this field of research, we require a fully annotated, digital 3D SC reference atlas for interpretation and comparison of data in mouse, the most widely used model system to study spinal circuit formation, somatosensory and motor behavior, axon regeneration and SC injury.

Here, we present accessible tools for efficient analysis of labeled cells and projections in whole mouse SC in the context of a novel 3D anatomical atlas that provides a common spatial framework for all studies. We have developed methods for oriented and parallel embedding of serial tissue pieces of the entire SC within a single block for controlled, synchronous cryo-sectioning and for automated imaging of sections. We have developed software, “SpinalJ”, to sort and register section images and to map the reconstructed data to a prototype 3D reference atlas. SpinalJ further combines tools for manual and automated analysis of identified cells and projections, and for data visualization. As an open resource, SpinalJ provides the community with high-throughput comparative analyses of neurons and their projections in whole SC.

## Results

To achieve a better understanding of the SC and functional spinal circuits, it is important to study the spatial relations of cells and projections in the context of the full SC. Moreover, comparison of data from individual animals and different labs requires a common coordinate framework for analysis. We have developed tools to accomplish this.

### Tools for efficient embedding and sectioning of whole spinal cord

Sectioning and imaging the entire adult mouse SC one section at a time while maintaining r-c order is challenging and time consuming. For efficient *in toto* immunohistochemical (IHC) or other label-based analysis of the SC, we first sought to reduce sectioning and processing time and to automate image acquisition.

#### Dividing SC tissue for synchronous and ordered sectioning

With a length of ~30mm, sectioning the cervical to lumbar region of adult mouse SC at 25μm produces ~1200 tissue sections. To reduce sectioning effort, the tissue can be divided into several consecutive pieces and embedded in parallel in the same block for synchronous sectioning. As the number of sections to collect (here called ‘block sections’) decreases by 1/N with the number of parallel embedded tissue pieces (N), maximizing N would reduce overall sectioning time. However, physical damage to the tissue is sustained with each cut at tissue piece boundaries. To balance sectioning speed and tissue preservation, we chose to cut the SC into nine tissue pieces of equal length (Fig.1A). For this, the fixed, cryo-protected SC was first cut into three equal pieces that together cover the cervical to lumbar extent of SC (Fig.1B). Each of these pieces was then further divided equally into three (Fig.1C), resulting in a total of nine tissue pieces (Fig.1D). This approach reduced the number of block sections and offered dense, regular spacing of tissue sections on the slide, saving time and materials for subsequent IHC processing and allowing for automated imaging.

**Figure 1:**
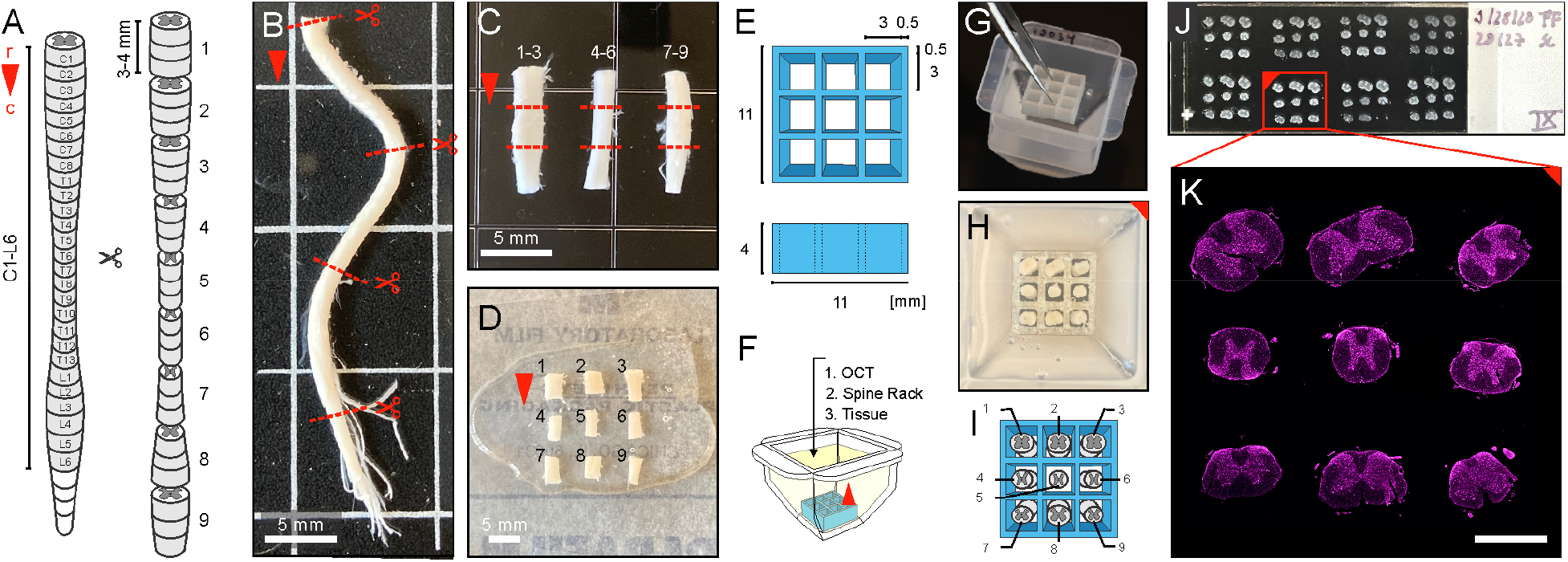
Embedding and sectioning of mouse spinal cord using SpineRacks. **A**) Cutting scheme for synchronous sectioning of nine parallel embedded tissue pieces of the adult mouse spinal cord. Red arrowhead indicates r-c orientation (panels A-D, F). **B**) The SC was divided into three equal pieces. Red dashed lines in B and C indicate cutting points. **C**) Each piece was then split again into three equal sized pieces, resulting in a total of nine pieces of SC tissue (D). **E**) Design and dimensions of SpineRacks. **F**) For tissue embedding, a SpineRack was sunk into a plastic mold filled with OCT embedding medium (g, see also Fig.S1) and tissue pieces were placed into the wells of the rack, each with its rostral end facing the bottom of the mold (H). Red filled image corner indicates the orientation of the tissue block for tracking as in J and K. **I**) Order of tissue pieces within the SpineRack. Pieces 1 to 3 were embedded left to right in the top row, pieces 4 to 6 in the middle row and pieces 7 to 9 in the bottom row. **J**) Eight cryostat block sections were collected in two rows (1-4, top left to right; 5-8, bottom left to right) on each slide (shown here as brightfield photograph). In this arrangement >1000 sections of adult mouse SC fit on 16 slides. **K**) Each block section, comprising nine tissue sections, was scanned on a slide scanning microscope and saved as a single image file (here shown labeled with Neurotrace 640/660 signal).

#### SpineRacks

Embedding the whole SC as several small tissue pieces in a regular array in a single block requires a structured approach to track tissue piece identity and to maintain r-c orientation. To facilitate fast, accurate and reproducible arrangement of the nine SC tissue pieces in an upright orientation, we developed 3D-printed, water-soluble polyvinyl alcohol (PVA) support scaffolds, here termed SpineRacks. We designed SpineRacks to fit into 12mm embedding molds and offer nine, 3mm by 3mm wide and 4mm deep wells arranged in a 3×3 grid (Fig.1E). After immersion of the SpineRack in a mold filled with embedding medium, SC tissue pieces were easily guided into the wells using blunt forceps (Fig.1F-I). The widest cervical segments fitted the wells in a diagonal orientation. The walls of each well prevented segments from falling over during embedding and block freezing. The SpineRack walls also acted as flow barriers, preventing already placed segments from drifting in the viscous embedding medium as other segments were being placed.

We chose PVA for production of SpineRacks, as this material is soluble in water and would be expected to dissolve slowly in water-based embedding media like OCT, offering two major advantages. First, while providing mechanical support to hold embedded tissue pieces in their orientation, the structure could be partially dissolved and softened for smooth sectioning during incubation in embedding medium before freezing. OCT contains 10% PVA and we argued that the shared material properties between SpineRack and surrounding embedding media would establish a continuous matrix, further promoting smooth sectioning. Second, after sectioning, SpineRack and OCT material surrounding the tissue sections could be washed away in PBS, eliminating any support components that could negatively affect staining protocols or contribute to background signals. Indeed, as predicted, SpineRacks dissolved in the OCT (Fig.S1) and the blocks sectioned smoothly, resulting in good tissue integrity. Moreover, during the first washes of histology protocols all embedding material washed away.

Using SpineRacks, sectioning effort was reduced almost 10-fold: all ~1200 sections from cervical, thoracic and lumbar levels of one adult mouse SC were collected in only ~130 block sections (each containing nine individual transverse sections of spinal tissue) on ~16 slides (Fig.1J,K). Moreover, the precise arrangement of tissue sections on the slide permitted automating acquisition, ordering and registration of images (see next section) such that processing of all sections can be achieved semi-automatically within a relatively short time (Table 1).

**Table 1:** Timing of whole spinal cord analysis

| Task | Hands on time | Unattended time | Total duration |
| --- | --- | --- | --- |
| Tissue embedding | 10-15 min | 20-30 min | 30-45 min |
| Cryo-sectioning | 1 h | - | 1 h |
| Immunohistochemistry | 1 h | Incubation for primary and secondary ABs | 24 h |
| Imaging (Slide Scanner) | 5-10 min | 3-5 h depending on the number of channels and resolution | 3-5 h |
| Image pre-processing (segmentation, cleaning, horizontal alignment, reformatting) | 15-30 min | 1-2 h depending on the number of channels | 1.5-2.5 h |
| Training Ilastik for machine learning based mesoscale mapping / cell detection | 15-30 min depending on the number of channels | - | 15-30 min |
| Image registration, atlas mapping and analysis | 1-5 min | 1-10 h depending on dataset size and the number of channels selected for Ilastik pixel classification | 1-10 h |
| Total time | 3-4 h | 28-40 h | 31-44 h |

Whilst our focus here is on the reconstruction of the cervical to lumbar SC of a single animal, SpineRacks can also be used to process in parallel one chosen smaller tissue region of interest (e.g. L3-L6) from nine different animals. Alternatively, SpineRacks with similar or adjusted geometries can be used for efficient and oriented sectioning of a variety of other tissues that are difficult to embed because of their low width-to-length ratio, such as muscle, organs or whole organisms like fish and insect larvae, or of multiple oriented samples of a given tissue. We have demonstrated such alternative applications of SpineRacks for parallel sectioning of multiple fish brains (Fig.S2) and, using Racks with larger wells, for sectioning precisely oriented adult mouse eyes (Fig.S3).

### Reconstruction of 3D image data from sections

The generation of ordered arrays of >1000 SC sections using SpineRacks demands computer assisted organization of images. To process whole SC image data, we have developed SpinalJ, a convenient plugin for ImageJ, which combines a series of software tools, together creating a seamless open source pipeline to facilitate image registration, atlas mapping and 3D analysis of SC sections. SpinalJ has been optimized, but is not limited, to work in combination with SpineRacks.

SpinalJ was conceived with reference to our previously published toolbox for the registration and analysis of brain sections, BrainJ (Botta et al., 2020), but with the additional development of a new preprocessing workflow to process section arrays, a novel 3D SC atlas and additional mapping and analysis options.

#### Image pre-processing in SpinalJ

For best flexibility and to accommodate various formats of input data, the image pre-processing workflow in SpinalJ has been organized in modules that can be executed independently. These comprise i) segmentation of block section images into individual tissue section images (Fig.2A,B), ii) compensation for lost tissue sections, iii) d-v re-orientation of sections, iv) r-c sorting of images (Fig.2C) and deleting/replacing damaged and out of focus images, as well as v) automated centering and horizontal alignment of SC sections (Fig.2D-F).

i. Utilizing the ordered array of SC tissue embedded in SpineRacks, SpinalJ automatically segments block section images that contain nine tissue sections into individual section images. Block section images that contain less than nine tissue sections cannot be segmented automatically and require simple manual placement of a segmentation mask on a preview displayed in SpinalJ.
ii. To maintain continuity within the 3D space of whole SC, SpinalJ uses the positional information of a list of sections that were lost during sectioning or washing to replace them by duplicating neighboring sections, thereby compensating for gaps in the image data (in our hands <15% of total images).
iii. Controlling d-v orientation of SC tissue pieces during embedding can be challenging, especially for thoracic segments. SpinalJ automatically aligns sections horizontally (see next section), but for this, approximate d-v orientation is required. If this is not the case, it is necessary to re-orient all sections from that tissue piece. SpinalJ creates a preview of all section images sorted by piece and the user can manually select pieces for re-orientation.
iv. For r-c sorting of images, SpinalJ uses either the alphanumeric order of image filenames or stage coordinate information extracted from the image metadata. For successful registration, the image dataset needs to be cleaned: occasionally, sections are damaged or out of focus and minor differences in the length of the tissue pieces may result in empty images at the beginning or end of the block. To correct for this, SpinalJ displays a preview of each section and the user selects whether to keep (intact image), replace with the neighboring section (damaged/out of focus image; or to delete (empty image) the current image.
v. Finally, horizontal alignment of SC sections facilitates the registration process leading to the final smooth continuity of the 3D reconstruction. We found that classical approaches to achieve horizontal alignment, like ellipse fitting (Pratt, 1987), did not perform well because SC sections differ dramatically in shape along the r-c axis (from highly elliptic to near round). Instead, we developed a method based on the bilateral symmetry of the gray matter stained using fluorescent Nissl stain Neurotrace (NT), (Quinn et al., 1995) or DAPI to determine the orientation of sections. For this, sections were centered and the NT/DAPI channel of images was thresholded and smoothened by Gaussian filtering. The image was then split vertically and the left half (L) was mirrored to match the orientation of the right half (R) and we calculated the average intensity of the absolute difference of both images (|L-R|, Fig.2D, shown for NT). This value is minimized when both image halves are mirror symmetrical, indicating horizontal orientation (Fig.2E). To determine the horizontal orientation for each section automatically, SpinalJ rotates section images by increments of 10° from −60° to +60° (covering the range of typical embedding orientations) and the angle at which the mean difference intensity is lowest is used to rotate the section (Fig.2F). Note that the distribution of mean difference intensities across all rotation angles has additional minima at ±90° and ±180° orientation. To ensure proper d-v alignment, sections oriented >90° and <-90° (upside down) were coarsely aligned, manually, first (see step iii).

**Figure 2:**
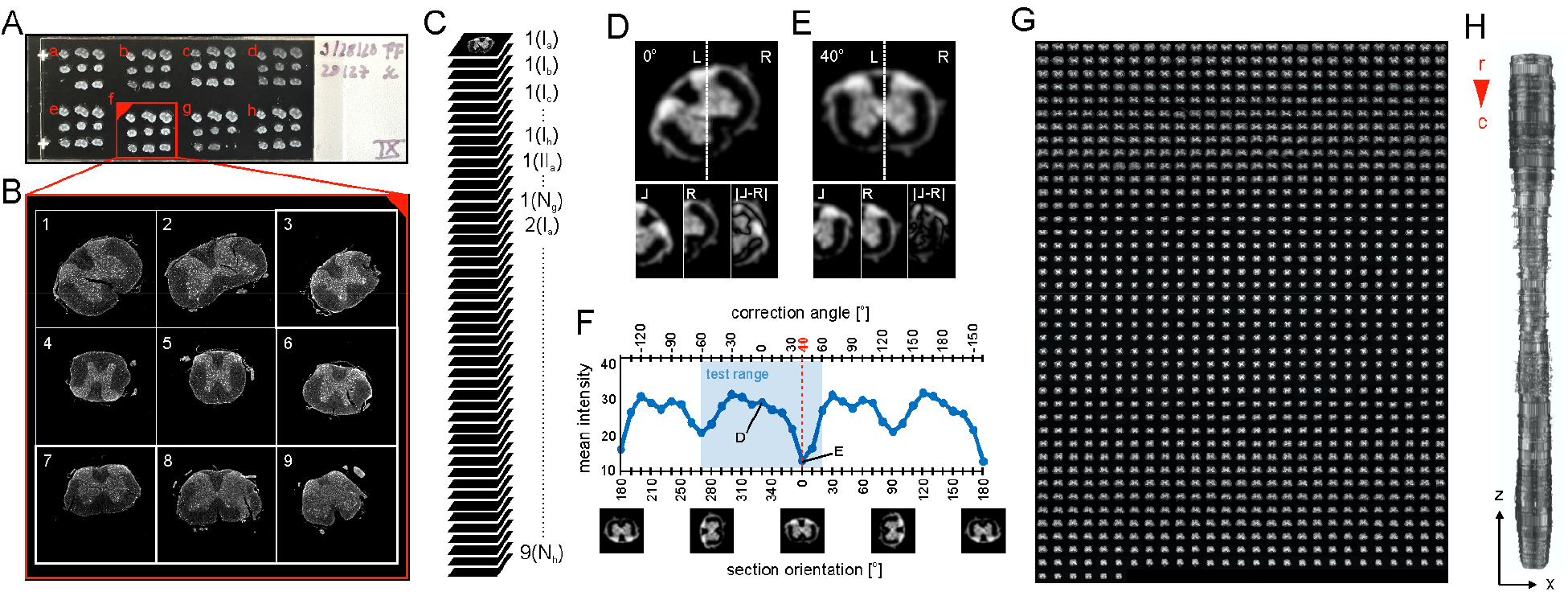
Image pre-processing and registration using SpinalJ. **A**) Slides were imaged on a slide scanning microscope (shown as brightfield photograph) and each block section (a-h) was saved as a single image file. The red filled corner indicates block section orientation (as in Fig.1H). SpinalJ then ordered images of block sections according to their position on the slide (a-h) to match sectioning sequence. **B**) Ordered images were split into nine individual tissue section images (1-9) (shown labeled with Neurotrace). **C**) Tissue section images were then sorted rostro-caudally in SpinalJ (section 1, slide I, block a = 1(Ia)). **D**) For horizontal alignment of sections, images were centered and thresholded using the Neurotrace or DAPI (not shown) channel. The resulting image was split vertically (dotted line) and the left half (L) was mirrored and overlaid with the right half (R) to calculate the average intensity of the absolute difference of both images (|L-R|). This value was minimized when both image halves aligned perfectly, indicating horizontal orientation (as illustrated in **E**). **F**) To determine horizontal orientation for each section automatically, images were rotated by increments of 10° from −60° to +60° (reflecting the range of typical embedding orientations; blue shaded area) and the angle resulting in the lowest mean difference intensity was applied to align the section. **G**) Montage of 1086 sorted and aligned section images of a SC sample spanning C1-S1. **H**) Dorsal view of the 3D reconstructed dataset (Neurotrace channel) shown in G after section registration.

#### Section Registration in SpinalJ

Image pre-processing in SpinalJ produces a continuous image stack of consecutive, intact tissue sections in r-c order (Fig.2G). To reconstruct a 3D dataset from these sections, a rigid body registration that preserves the shape of the tissue sections (Thévenaz et al., 1998) was performed on a contrast-enhanced and down-sampled (10μm/pixel) copy of the registration channel (DAPI and NT were both found to be suitable for this purpose). The re-scaled transformations were subsequently applied to all channels at the desired resolution for analysis (typically 2μm/pixel), which yielded a registered 3D SC volume (Fig.2H).

### Creation of a 3D reference atlas for mouse SC

Comparative analysis and interpretation of whole SC data from registered SC sections requires a standard framework like a reference atlas, which, to date, was unavailable. We have therefore built a 3D atlas for mouse SC. For this we used, as a base, the 34 annotated 20μm sections (one for each spinal segment) of the Allen Spinal Cord Atlas (Allen Institute for Brain Science, 2008) (Fig.3A,B). First, we digitized a total of 73 regions within the Allen Institute dataset using an intensity map (Fig.3C). These annotated section images, and their corresponding Nissl images, were then manually edited to be symmetrical and free from idiosyncratic features or damage that may impair their use as a template for registration. Next, we used the Nissl images to generate a 3D volume. For this, images were first processed using attenuation correction to compensate for section-to-section intensity variations and were then aligned using rigid body registration (Thévenaz et al., 1998). Images typifying each SC segment were then stacked with a spacing of 20μm to fill the average length of each spinal segment, thereby generating a 3D volume representing the entire SC (Fig.3D). Segment lengths and the position of segment boundaries were determined, informed by the relative positions of sections chosen by the Allen Institute for annotation and by the relative lengths of segments as reported in the literature (Harrison et al., 2013) (Fig3.F). This 3D template serves as a common framework and was used to align all experimental datasets. To complete the atlas, the transformation parameters obtained from registering Nissl sections were applied to the digital annotation images (Fig.3E). For convenient analysis in SpinalJ, we created additional atlas region groups, combining individual annotated regions into relevant quantification clusters (e.g. region group ‘Lamina V’ combines atlas regions 5Sp, 5SpL, 5SpM, D, SDCom, CeCv and IMM5). These calculated region groups allow SpinalJ users to evaluate data at different levels of anatomical detail. A list of all atlas regions and region groups is provided in Table 2.

**Figure 3:**
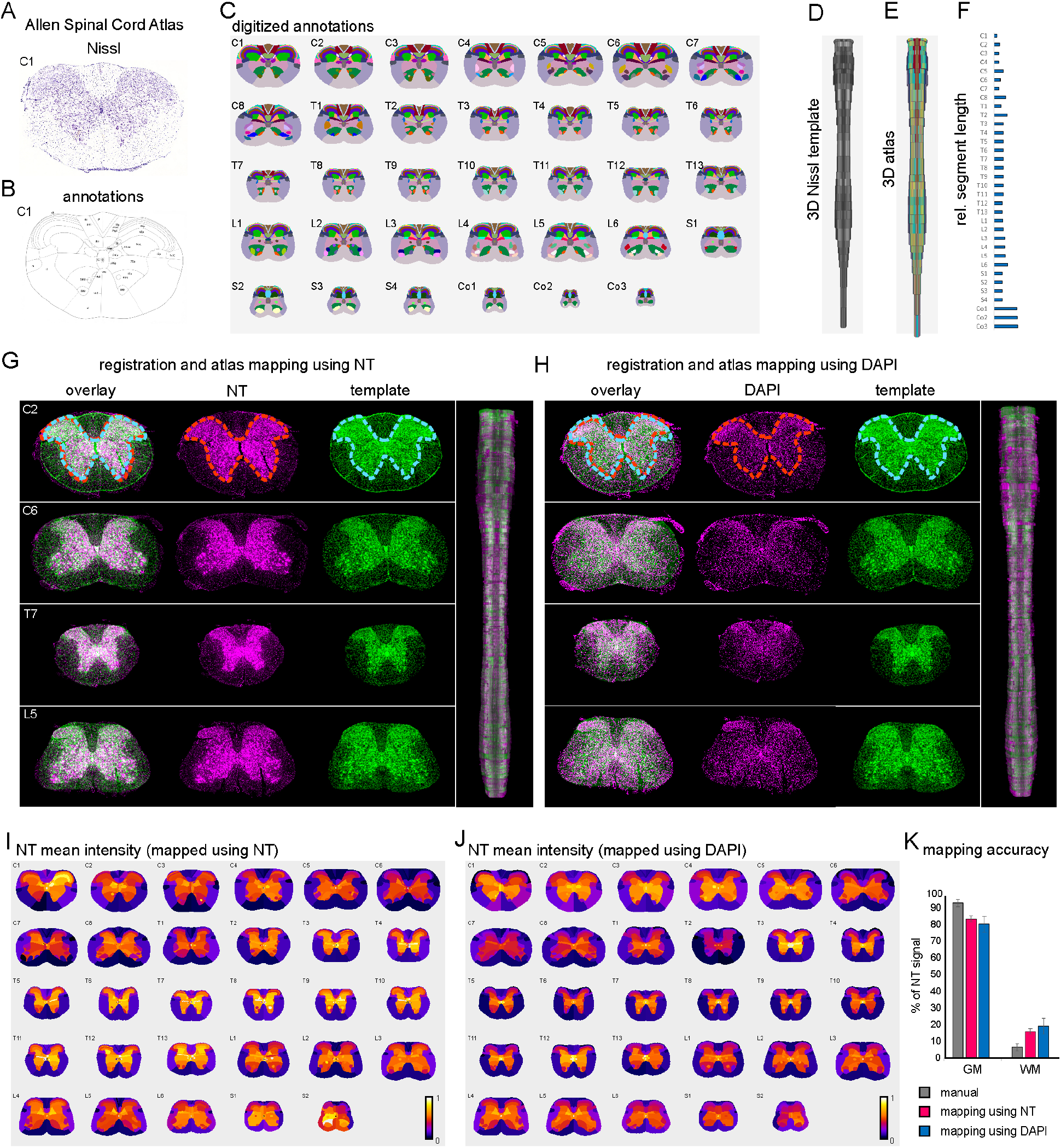
Mapping registered spinal cord sections to a novel 3D atlas in SpinalJ. Creation of the 3D reference atlas used 34 Nissl-stained sections (**A**) (shown for C1) and corresponding annotated sections (**B**) of the Allen Institute Spinal Cord Atlas (Allen Institute for Brain Science, 2008). **C**) Annotations in all sections (1 for each spinal segment, C1-Co3) were translated into a grayscale pixel code (here visualized in color). **D**) A 3D Nissl template (dorsal view) was created by registering and extruding the 34 Nissl sections representing each spinal segment. **E**) The transformation to create D was then applied to the annotated sections (C) to generate a 3D annotated atlas. **F**) Segment boundaries were placed according to the relative positions of sections in the Allen Institute’s dataset and the relative lengths of spinal segments (blue bars) as reported in the literature (Harrison et al., 2013). **G**) Example of mapping the same experimental dataset (magenta) to the Nissl template (green) using either NT or **H**) DAPI. For ease of comparison, dashed lines are presented in G and H to indicate the boundary between gray matter (GM) and white matter (WM) in C2. **I**) Heatmaps of mean NT intensities per atlas region after template mapping with NT in SpinalJ, summarized for each spinal segment. **J**) Heatmaps of mean NT intensities per atlas region after template mapping with DAPI in SpinalJ, summarized for each spinal segment. **K**) Distribution of NT signal between GM and WM along the entire SC of four animals, measured manually (gray bar) and after SpinalJ atlas mapping using NT (magenta bar) or DAPI (blue bar) counterstaining. NT signal in GM: 93.68% (manual), 84.09% (±1.73) (NT), 80.92% (±4.76) (DAPI). NT signal in WM: 6.32% (manual), 15.91% (±1.73) (NT), 19.08% (±4.76) (DAPI). Error bars represent standard deviation of the mean. With reference to manually determined signal distributions, SpinalJ mapping accuracy was determined 89.76% (NT) and 86.37% (DAPI), respectively.

**Table 2:**
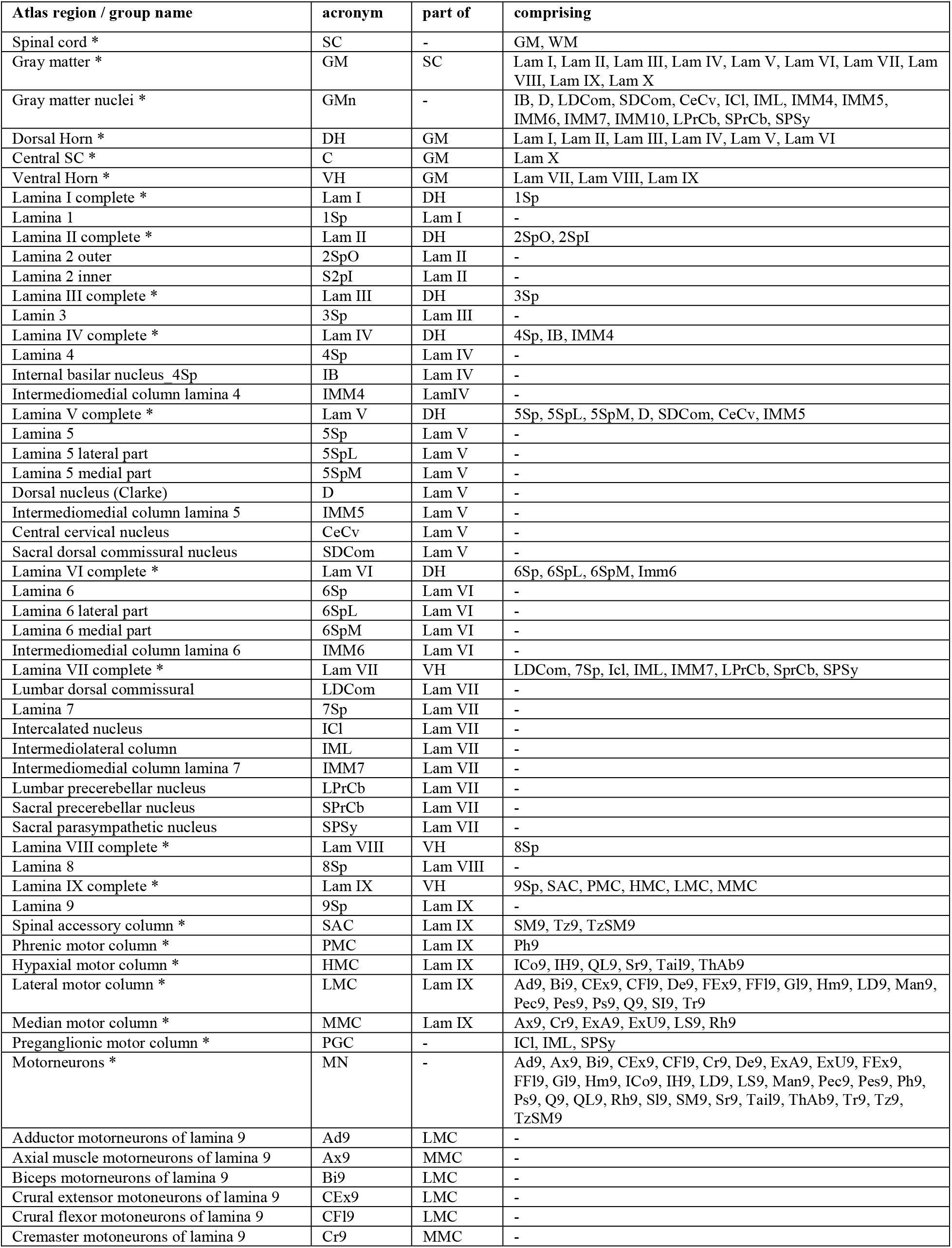
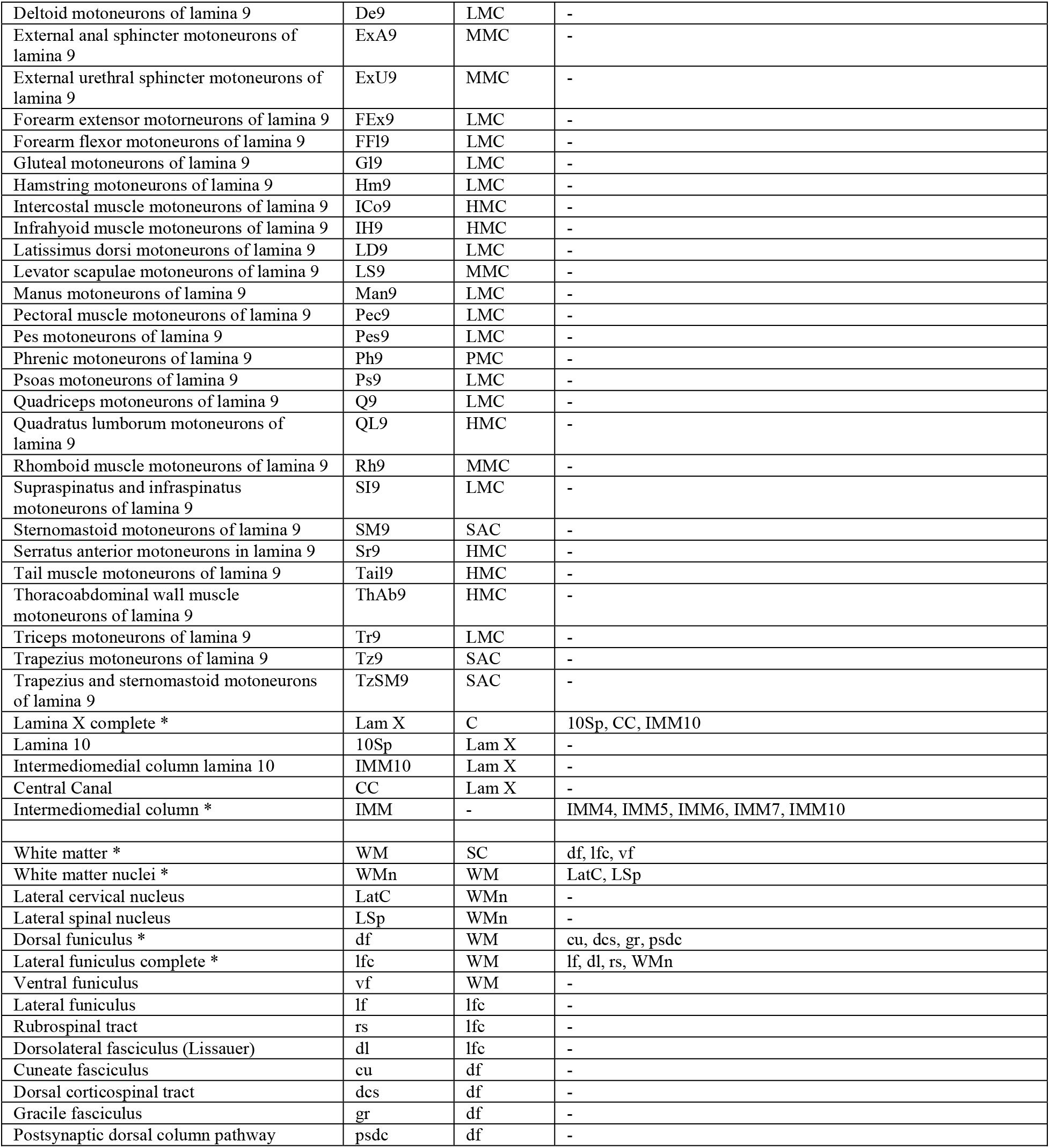
Atlas regions and calculated region groups (* marks calculated region groups)

### Using SpinalJ for atlas mapping

To bring experimental data into anatomical context, we programmed SpinalJ to map the registered sections to the 3D Nissl template and overlay them with the 3D annotations for analysis. We refer to this process as ‘atlas mapping’. For this, the registration channel of the experimental data was first resampled to match the resolution of the atlas template and the r-c segment range of the experimental dataset. Registration of the resampled data to the relevant portion of the atlas template was performed using Elastix (Klein et al., 2010). Briefly, a 3D affine transformation followed by a 3D B-spline transformation was performed with Mattes Mutual Information used to calculate similarity.

To validate atlas mapping using NT and DAPI for registration of sections and mapping to the Nissl template, we inspected the alignment of gray matter (GM) and white matter (WM) regions between experimental data and template. Visual inspection using either marker showed good alignment (Fig.3G,H). To assess the mapping accuracy using either marker quantitatively, we first outlined the gray and WM in 39 randomly selected sections from four animals by hand and determined the mean intensities of thresholded NT signal in each region using Fiji. NT binds the ribosomal RNA associated with the rough endoplasmic reticulum in the soma and dendrites of neurons (Quinn et al., 1995) and signal should thus be restricted to the GM. Indeed, we found strong signal in the GM of SC sections stained with NT660 (93.68% of total NT signal, N=4, n=39), and only sparse, punctate signal in the WM (6.32% of total NT signal) (Fig.3K). We then compared these manually determined values to the signal distribution within the annotated GM and WM regions after atlas mapping using SpinalJ. We found that, across all spinal segments, 84% of NT signal was mapped to the GM and 16% to the WM, using NT for section registration and mapping (Fig.3I,K), indicating a mapping accuracy of 90%. Mapping was slightly less accurate using DAPI (86%), with 81% of NT signal in the GM and 19% in the WM (Fig.3J,K). These results confirm that SpinalJ offers highly accurate template mapping of 3D reconstructed sections.

### Analysis of cells and projections using SpinalJ

For the analysis of position and connectivity of neurons within the atlas, SpinalJ offers options to detect cells and projections automatically, via image segmentation, but can also import a list of manually determined coordinates. Options for automatic segmentation can be chosen based on needs for processing time and detail of analysis. The quickest and simplest segmentation method, ‘binary thresholding’, applies a user-specified intensity threshold to separate high-from low-intensity signal. To tune segmentation for cell detection, ‘find maxima’ isolates the positions of local intensity maxima around a user specified intensity value. Most detailed segmentation can be achieved using ‘machine learning segmentation’ for detection of both cells and neurites. This is based on distinct pixel probabilities derived from training pixel classifiers for features of interest using Ilastik (Berg et al., 2019). All approaches allow for additional filtering based on intensity and object size. Extracted spatial information of cells and projections is then transformed to the SC atlas space, and a reverse mapping approach is used to measure cell and projection density within annotated regions. SpinalJ outputs these data in both table and image formats for easy exploration of the data.

To test the mapping accuracy of SpinalJ, we analyzed labeled neurons with well characterized, spatially defined distributions within sub-regions of the SC gray and WM.

#### Mapping of neuronal position

As a first validation set, we labeled SC sections with an antibody against choline acetyltransferase (ChAT), a marker for cholinergic neurons (Fig.4A). The distribution of ChAT immunoreactivity in adult SC has been described; it is restricted to motor neurons (MNs) of lamina IX, neurons in the intermediolateral column, the intercalated nucleus, the sacral parasympathetic nucleus and medial part of lamina VII, as well as in the central autonomic area and the central canal cluster neurons of lamina X (Barber et al., 1984; Heise and Kayalioglu, 2009). To assess the overall distribution of ChAT signal after mapping in SpinalJ, we determined the mean intensity of ChAT signal per atlas region for each spinal segment. We then visualized the mapped intensity data as an intensity heatmap montage to assess the signal distribution across all segments (Fig.4B). This analysis demonstrated that signals map predominately to the ventral GM regions of SC. To visualize the data within larger region groups across segments, we generated heatmap matrix plots, which showed highest intensities in laminae VII, IX, X and also VIII (Fig.4C). Additionally, lower intensity signals were mapped to more dorsal laminae, where visual inspection revealed background immunofluorescence but no ChAT^+^ cells, highlighting the importance of a good signal-to-noise ratio for clean interpretation of data. While mapping mean intensities is the fastest way to analyze data in SpinalJ, we applied alternative segmentation methods to refine the data.

**Figure 4:**
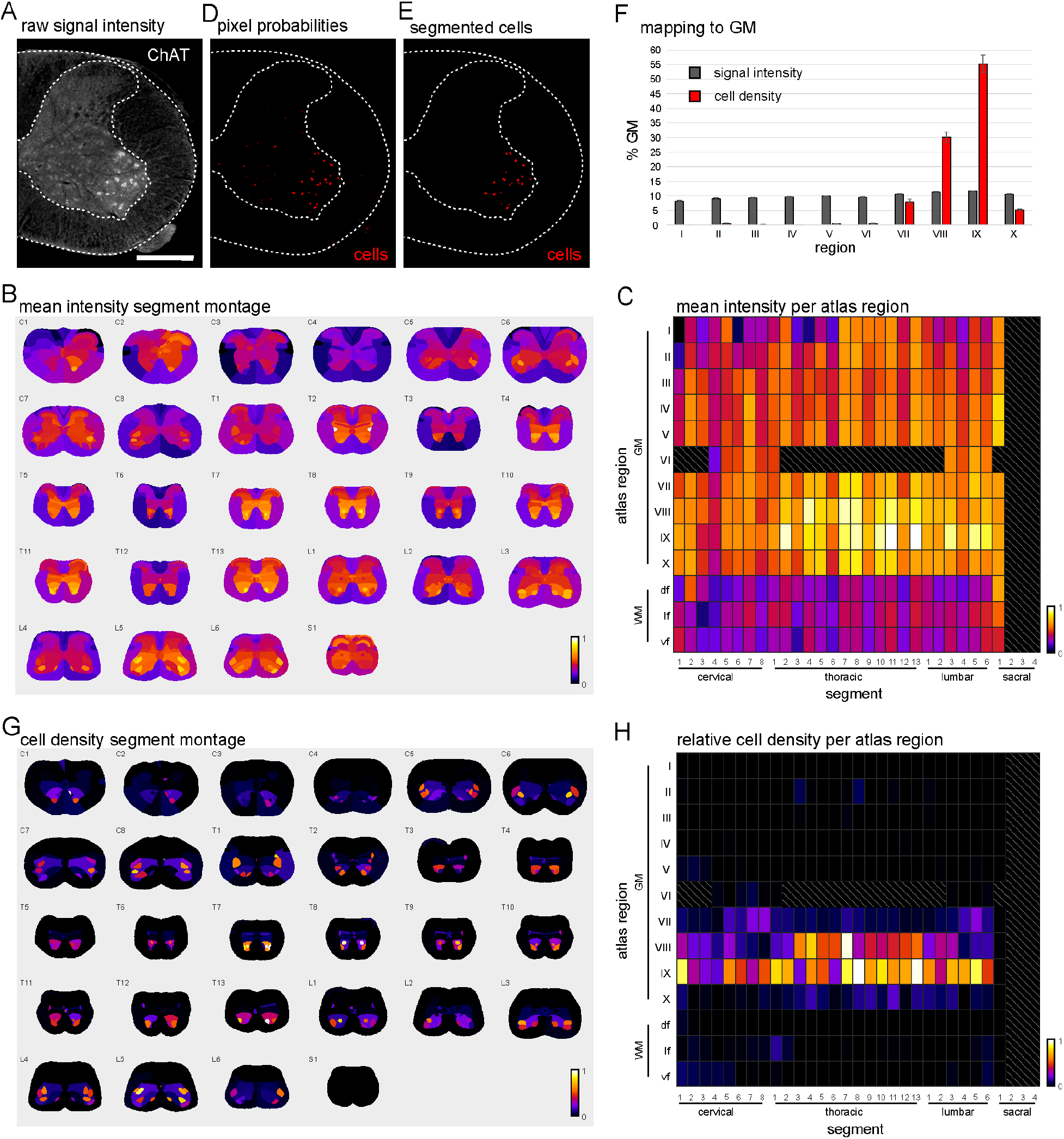
Intensity and cell density mapping of ChAT^+^ neurons. **A**) Hemisegment of SC section labeled with ChAT antibody. Scale: 500μm. **B**) Heatmap montage showing the spatial distribution of mean ChAT intensity per atlas region and segment. **C**) Heatmap matrix plot showing the distribution of mean ChAT intensity in atlas region groups of the gray (laminae I-X) and white matter (df: dorsal funiculus, lf: lateral funiculus, vf: ventral funiculus). **D**) Pixel probabilities for classifiers ‘cell’ (red) after training in Ilastik. **E**) Cells detected after image segmentation using pixel probabilities. **F**) Relative distribution of ChAT signal intensities (gray bars) and cell densities (red bars) within atlas regions of the gray matter (GM) (laminae I-X). Error bars indicate standard deviation between values from both hemisegments. **G**) Spatial distribution of relative ChAT^+^ cell density per atlas region and segment. **H**) Heatmap matrix plot showing the distribution of relative ChAT^+^ cell density in atlas region groups. Gray hatched areas in C and H mark regions without data.

In order to extract features of interest and to eliminate background signal, we used automatic image segmentation via Ilastik machine learning pixel classification (Fig.4D,E). After initial training, segmentation parameters can be applied to multiple datasets for high-throughput analysis. We validated the accuracy of the automatic cell detection by visually inspecting the overlay of detected cells and ChAT signal provided by SpinalJ (Fig4A,E). Quantification of a randomly selected subset of 20 sections confirmed that 89% of labeled cells were detected using Ilastik, attesting to the accuracy of this segmentation approach.

The coordinates of identified cells were then used to calculate cell densities per atlas region (Fig.4G,H). This approach provided a much clearer distribution of ChAT^+^ cells within the SC than intensity mapping (Fig.4F) and reproduced the known distribution with the exception of cells mapped to lamina VIII (potentially as a result of imprecise lamina IX annotations; see discussion). Cell densities were greatly reduced in laminae I-VI and X, indicating that mean intensity mapping included non-cellular/background signal. These results illustrate the usefulness of mean intensity mapping for quick analysis of overall signal distribution that can be further refined using more detailed segmentation approaches, especially in the presence of high background.

#### Mapping of peripheral fiber terminals

Next, we tested the detection and mapping accuracy of sensory projections labeled by Isolectin B4 (IB4). IB4 binds non-peptidergic dorsal root ganglion (DRG) neurons and their afferents, which innervate primarily the inner part of lamina II (lamina IIi) and, to a lesser extent, lamina II outer (lamina IIo) (Molliver et al., 1995; Silverman and Kruger, 1990; Takazawa et al., 2017).

Mapping of IB4-FITC signal showed a concentration of mean intensities in the superficial dorsal horn with a trend towards laminae I and II, at all r-c levels (Fig.5A,B). However, while the highest intensities were observed in laminae I, II and III, lower but significant signal was also detected in the intermedioventral SC (laminae IV-X, Fig.5C), suggesting mapping of background signal. Analysis of projection densities per atlas region after machine learning image segmentation (Fig.5D,E) revealed that the highest projection densities were mapped to lamina II (Fig.5G,H). In contrast to mean intensities, projection densities in laminae I and III were significantly lower than in lamina II (reduced by 19.0% and 18.5%, respectively) and close to zero in laminae IV-X (Fig.5F). Within lamina II, we found slightly higher densities in lamina IIi compared to lamina IIo, although this trend was not statistically significant (Fig.5F, inset). Thus, SpinalJ also performs well in mapping axon terminals and is able to filter out signals of particular interest using segmentation.

**Figure 5:**
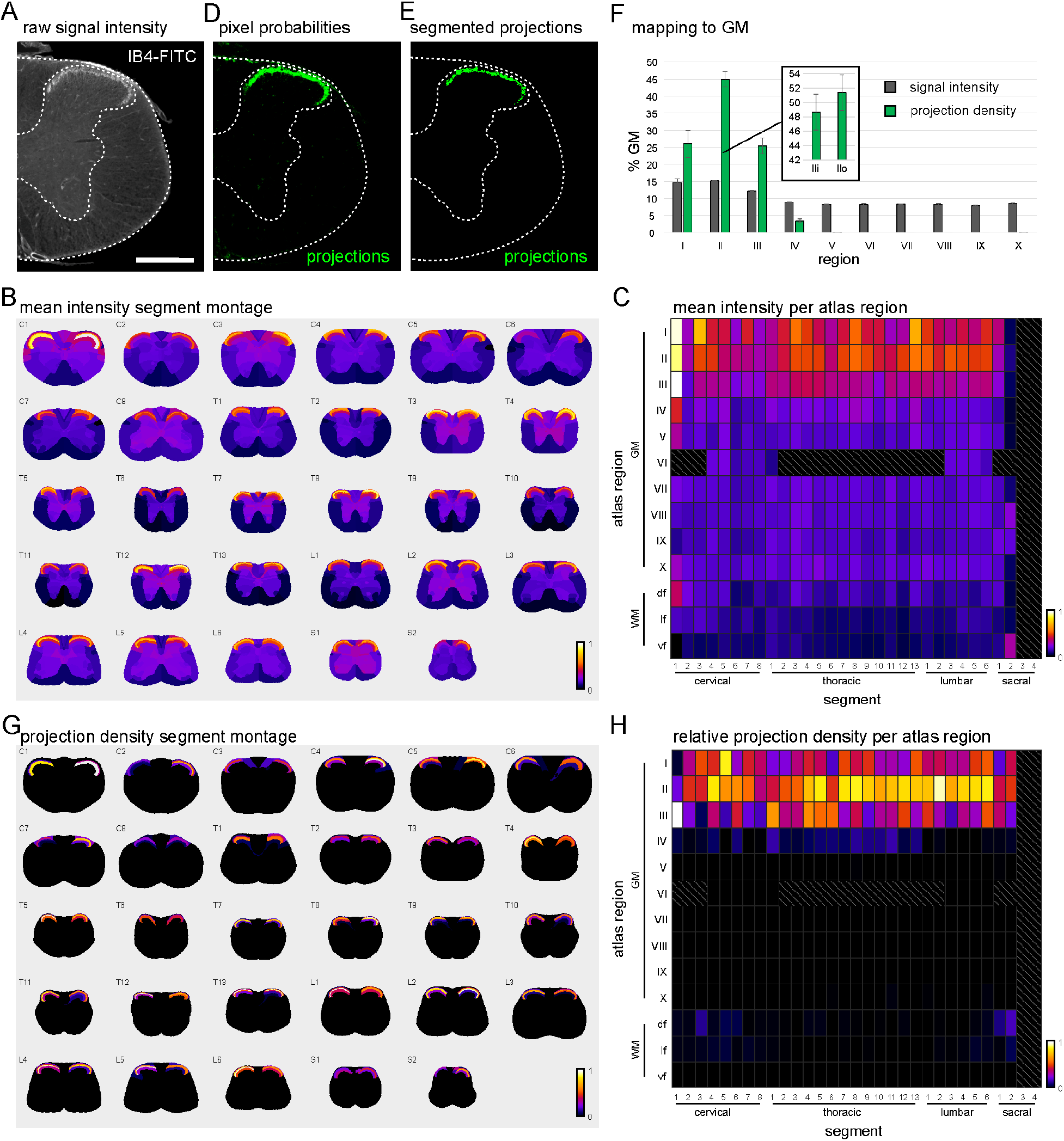
Intensity and projection density mapping of IB4^+^ fibers. **A**) Hemisegment of SC section labeled with IB4-FITC. Scale: 500μm. **B**) Spatial distribution of mean IB4 intensity per atlas region and segment. **C**) Distribution of mean IB4 intensity in atlas region groups of the GM (laminae I-X) and WM (df: dorsal funiculus, lf: lateral funiculus, vf: ventral funiculus). **D**) Pixel probabilities for classifiers ‘projections’ (green) after training in Ilastik. **E**) Projections detected after image segmentation using pixel probabilities. **F**) Relative distribution of IB4 signal intensities (gray) and projection densities (green) within atlas regions of the GM (laminae I-X). Inset shows projection densities within laminae II (IIi and IIo). Error bars indicate standard deviation between values from both hemisegments. **G**) Spatial distribution of relative projection density per atlas region and segment. **H**) Distribution of relative projection density in atlas region groups. Gray hatched areas in C and H mark regions without data.

#### Mapping of long-range projections

To test the ability of SpinalJ to map axonal projections over long r-c distances and seamlessly across multiple segments, we labeled corticospinal projection neurons and their axons forming the corticospinal tract (CST) by unilateral injection of AAV-tdTomato into the cortex of an adult mouse (Fig.6A,B). We determined the position of injection sites and found AAV infected, tdTomato^+^ neurons in the primary and secondary motor area (n=2367, 60% total labeled cells), primary somatosensory area (n=1022, 26%) and anterior cingulate area (n=513, 13%), with few cells (n=22, 1%) in non-CST regions (mainly CA2 and CA3) of the left hemisphere. Thus, 99% of labeled neurons were located within nuclei that contribute to CST.

**Figure 6:**
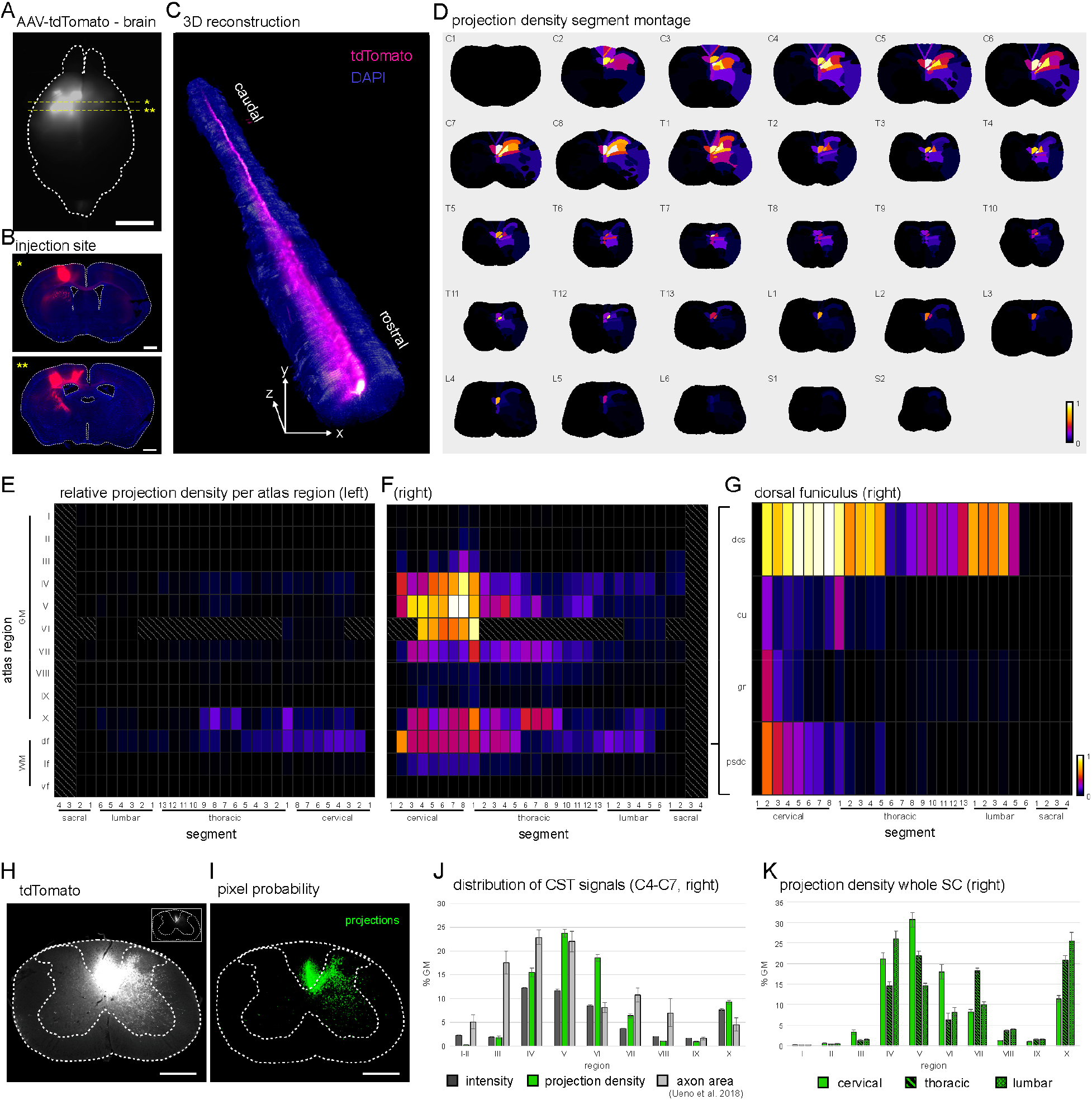
Intensity and projection density mapping of AAV labeled CST axons. **A**) tdTomato signal in a dorsal view of the brain shows AAV-tdTomato injection sites into left sensory and motor cortex. Yellow dotted lines indicate positions of coronal sections shown in B. Scale bar: 3mm. **B**) Coronal sections of the brain at levels indicated in a, showing a composite of tdTomato (red) and DAPI (blue) channels. Scale: 1mm. **C**) 3D reconstruction of registered SC sections. Image shows the 3D dataset (composite of tdTomato (magenta) and DAPI (blue) channels) in a frontal and dorsal view. **D**) Spatial distribution of relative projection density per atlas region and segment. **E**) Distribution of relative projection density in atlas region groups of the GM (laminae I-X) and WM (df: dorsal funiculus, lf: lateral funiculus, vf: ventral funiculus) within the left (ipsilateral) and right (**F**, contralateral) hemisegments. Gray hatched areas mark regions without data. **G**) Distribution of relative projection density in atlas regions within the dorsal funiculus of the right hemisegment (dcs: dorsal corticospinal tract, gr: gracile fasciculus, psdc: postsynaptic dorsal column pathway, cu: cuneate fasciculus). See also Fig.S4. **H**) Cervical SC section showing the distribution of tdTomato signal. Inset shows the same image reduced in brightness. Scale: 500μm. **I**) Pixel probabilities for classifiers ‘projections’ (green) after training in Ilastik. **J**) Relative distribution of mean intensities (dark gray bars) and projection densities (green bars) within atlas regions of the GM of segments C4-C7. Light gray bars show the relative distribution of CST axon area as measured in a manual mapping study by Ueno and colleagues (Ueno et al., 2018) for comparison. **K**) Distribution of relative projection densities within atlas regions of the GM of cervical (plain green bars), thoracic (diagonally banded bars) and lumbar (checkerboard patterned bars) segments. Error bars indicate standard deviation of values from all segments within the analyzed range.

In cervical SC, CST neurons have been reported to innervate laminae III-VIII and lamina X of the contralateral GM (Ueno et al., 2018). After section registration in SpinalJ, the 3D reconstruction of our data showed a continuous CST with lateral branches innervating the dorsal horn (Fig.6C). Analyzing the distribution of projection densities after segmentation of CST signals (Fig.6H,I), we found high densities in laminae IV, V and VI of the contralateral GM, with lower densities in laminae III, VII and X (Fig.6D-F). In the WM, the highest projection densities were found in the contralateral dorsal funiculus (df, Fig.6f) and, within the df, projection densities concentrated in the dorsal CST region (dcs, Fig.6G). Low projection densities were also identified in lamina X and df of the ipsilateral hemisegment. In all regions, projection densities declined along the r-c axis, with the highest densities in cervical segments, reflecting the thinning of the tract towards the caudal end of the SC. Mapping mean signal intensities showed overall similar signal distributions (Fig.6J).

Both of our analyses revealed signal distributions that closely matched the results of a manual mapping study on segments C4-C7 (Ueno et al., 2018) (Fig.6J, light gray bars), validating SpinalJ mapping. A major advantage of analysis of whole SC in SpinalJ is that the entire projection can be visualized and projection densities measured at all spinal levels simultaneously (Fig.6K). Moreover, the ability to segregate features such as bright axon bundles (Fig.S4A) and dimmer lateral branches (Fig.S4B) in the same image channel using Ilastik is a powerful option in SpinalJ for selective mapping of features of interest (Fig.S4C,D). This principle of using segmentation in SpinalJ can be applied also to discriminate morphologically distinct compartments of neurons. For example, soma and neurites can be analyzed separately in the same image channel, as demonstrated in Fig.S5A-E.

Thus, while tracing individual axons might be impracticable in registered sections due to slight registration offsets, the quick assessment of axonal tracts and terminations in 3D using SpinalJ provides a new approach to studying SC connectivity and regeneration of axons after injury.

### Mapping and comparing multiple samples using SpinalJ

#### Mapping accuracy of the same cell population across samples

The approach of mapping section data to a standardized template in principle permits the comparison of spatially discrete populations of neurons across different animals. To assess the alignment accuracies of multiple datasets, we mapped and compared the positions of ChAT^+^ neurons (Fig.7A) from three whole SCs. For this, we plotted the 3D cell positions, color coded for each animal (Fig.7B). The overall 3D distribution of this cell population was represented in each sample. To quantify inter animal mapping differences within each spinal segment, we projected the cell position data of each spinal segment along the r-c dimension, as shown in Fig.7C. We then calculated the center of mass (centroid position) of the transverse 2D cell distribution within each hemisegment and spinal level for each animal (Fig.7D,E). Next, we calculated and averaged the pairwise distances between all centroids as an indication of mapping precision. Between animals and across all spinal levels we found an average centroid distance (mapping disparity) of only ~13μm (Fig.7F). These results show that mapping data from different animals can be accurately achieved in SpinalJ and suggest that this tool is well suited to explore the distributions of novel markers.

**Fig 7:**
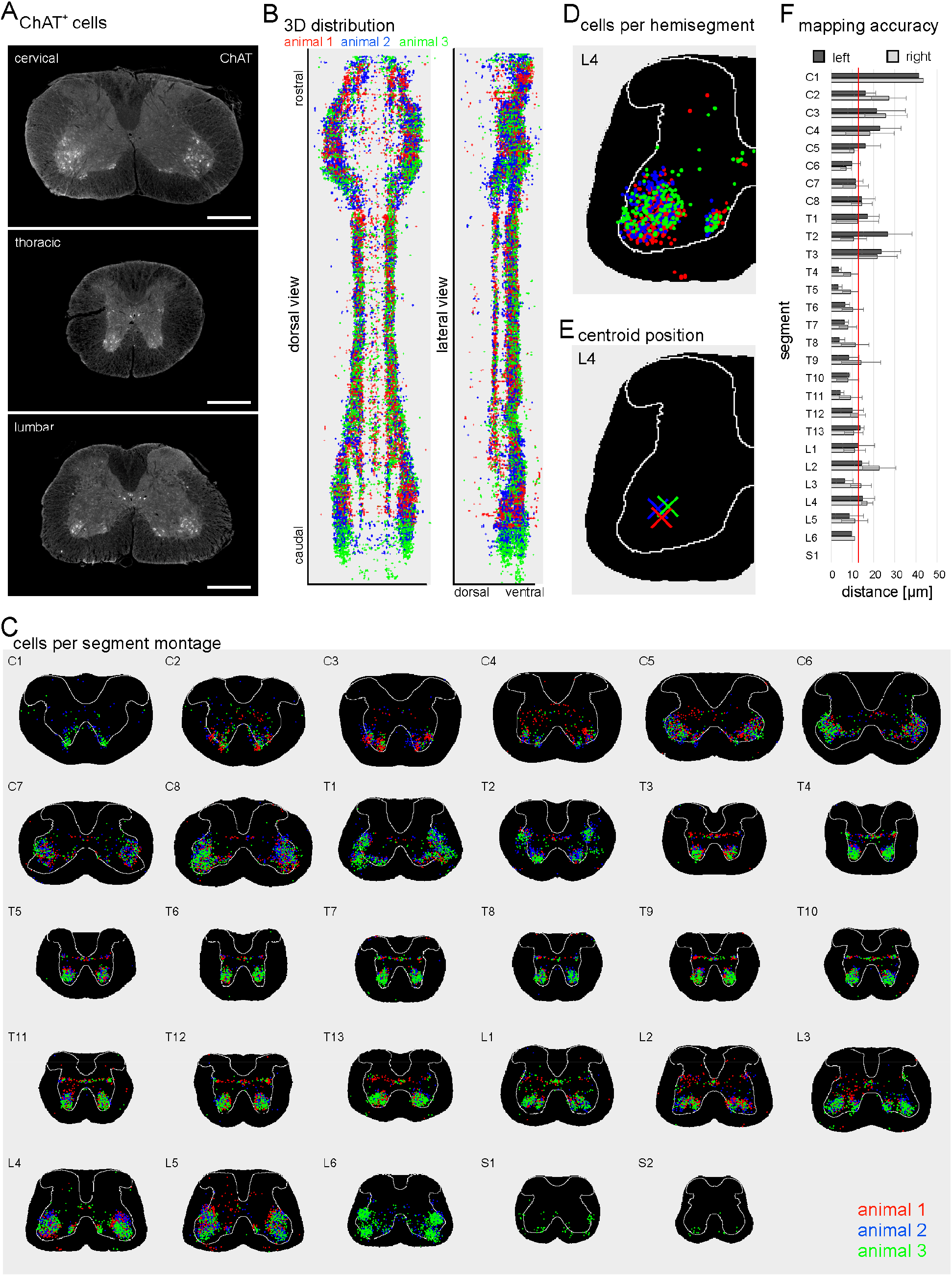
Mapping of ChAT cells in multiple animals. **A**) Sections labelled with anti-ChAT antibody. Scale: 500μm. **B**) 3D distribution of the positions of ChAT^+^ cells detected in three different samples (animals 1-3, red, blue, green) in a dorsal (left) and lateral (right) view. **C**) Spatial distribution of ChAT^+^ cells from different samples within each spinal segment. **D**) Cell distributions within each hemisegment (shown for L4, left hemisegment) were analyzed to determine the center of mass (centroid) for each animal, spinal level and hemisegment (shown for L4, left in **E**). **F**) Average pairwise centroid distances indicate the mapping offset between animals within the left (dark gray) and right (light gray) hemisegment at different spinal levels. Red line indicates average centroid distance across all segments (13.3μm). Error bars indicate standard deviation of the mean.

To test the ability of SpinalJ to delineate cell populations with unknown distributions, we chose to map the subset of MNs that derive exclusively from progenitors expressing the Forkhead domain transcription factor 1 (Foxp1). These neurons comprise the cholinergic neurons of the embryonic lateral motor column (LMC) and the preganglionic motor column (PGC) of lamina IX (Dasen et al., 2008; Morikawa et al., 2009), but they have not been mapped in the adult, since Foxp1 is also expressed in other spinal neurons, precluding selective labeling. To characterize the 3D distribution of cholinergic Foxp1 neurons in the adult, we used a new intersectional mouse line that expresses tdTomato in VAChT^+^, Foxp1^+^ neurons (Ng et al., in preparation).

Mapping VAChT^+^/Foxp1^+^ cells using automatic cell detection in SpinalJ revealed large clusters of cells in the ventrolateral SC at limb levels within the annotations of the LMC (Fig.8A,B, black arrowheads, Fig.S5). In each hemisegment, an additional, thinner cluster of cells was observed extending caudally from the cervical LMC and spanning the intermediolateral thoracic and upper lumbar segments as part of the PGC (Fig.8A,B, white arrowheads, Fig.S5). These findings were in line with the expected distributions of these cells (Fig.8C). In the embryo, LMC neurons have been identified only at limb levels, whereas PGC neurons were also found in thoracic and upper lumbar segments T1-L2 (Jessell, 2000; Prasad and Hollyday, 1991; Stifani, 2014; Tsuchida et al., 1994). Surprisingly, however, we also observed a smaller cluster of cells in the extreme ventral horn of thoracic segments within the annotations of the hypaxial (HMC) and median motor columns (MMC) (Fig.8A,B, yellow arrowheads, Fig.S5), regions not previously thought to include any cholinergic Foxp1 lineage neurons (Stifani, 2014). Minimal mapping offsets between samples (Fig.8B,D-F) suggest that these cells were not mismapped to these regions. Moreover, we confirmed that these cells are indeed cholinergic neurons by co-staining with ChAT antibody (Fig.9A-D), ruling out the possibility of unspecific labeling or segmentation artifacts. Thus, using SpinalJ, we identified a previously undescribed population of cholinergic Foxp1^+^ neurons, which require further characterization, and mapped the 3D distribution of LMC and PGC MNs.

**Figure 8:**
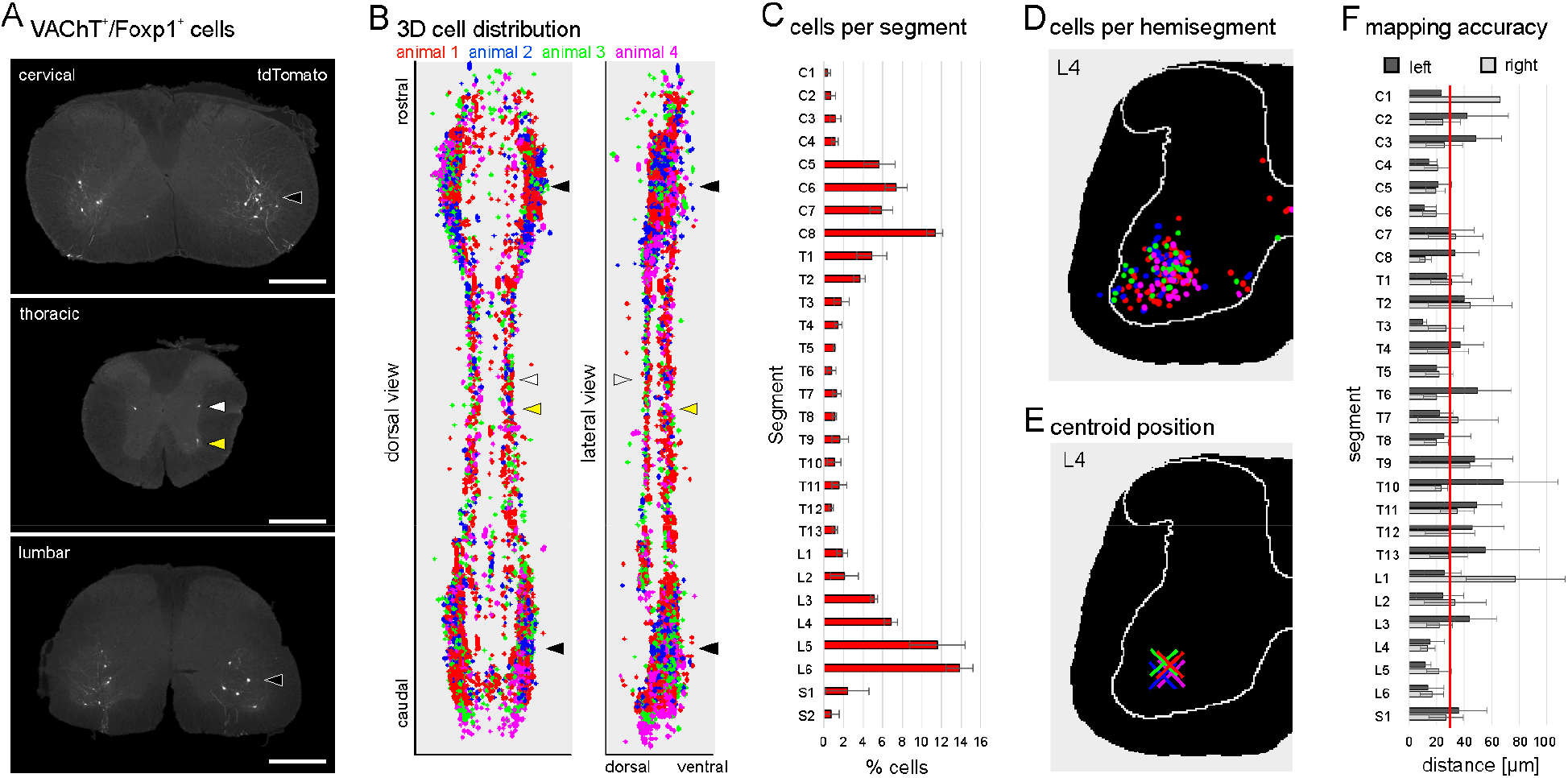
Spatial distribution of VAChT^+^/Foxp1^+^ cells. **A**) SC sections showing tdTomato^+^ cells. Arrowheads mark cell clusters in the lateral motor columns (LMC, black), preganglionic motor columns (PGC, white) and hypaxial/median motor columns (HMC/MMC, yellow). **B**) 3D distribution of the positions of VAChT^+^/Foxp1^+^ cells detected in four different samples (animals 1-4, red, blue, green, magenta) in a dorsal (left) and lateral (right) view. Arrowheads mark motor columns as in A. See Fig.S5 for 2D distributions of cells in individual segments. **C**) Average number of VAChT^+^/Foxp1^+^ cells per spinal segment across all four animals. Error bars represent the standard deviation between animals. **D**) Cell distributions within each hemisegment (shown for L4, left hemisegment) were analyzed to determine the center of mass (centroid) for each animal, spinal level and hemisegment (**E**). **F**) Average pairwise centroid distances indicate the mapping offset between animals within the left (dark gray) and right (light gray) hemisegment at different spinal levels. Red line indicates the average centroid distance across all segments (30.3μm). Error bars indicate standard deviation of the mean.

**Figure 9:**
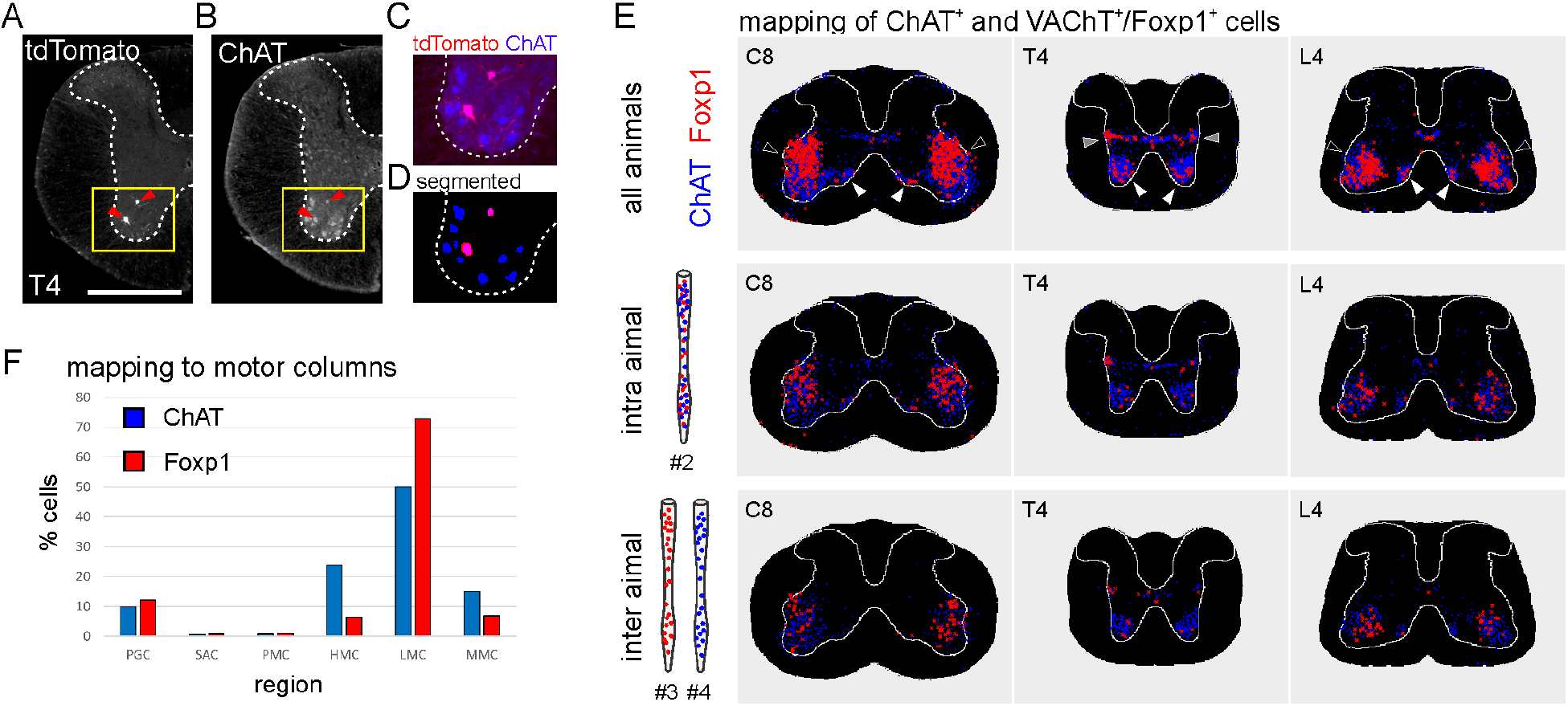
Inter animal mapping accuracy of VAChT^+^/Foxp1^+^ and ChAT^+^ cells. **A**) Section showing VAChT/Foxp1:tdTomato labeled neurons (arrowheads) in the ventral thoracic SC (T4). Scale: 500μm. **B**) ChAT counterstaining of section in A. **C**) Overlay of tdTomato (red) and ChAT (blue) signal from A and B. **D**) Overlay of segmented tdTomato (red) and ChAT (blue) signals used for cell detection in the same field of view. **E**) Top row: Overlay of the positions of detected VAChT^+^/Foxp1^+^ (Foxp1) cells (red) and ChAT^+^ cells (blue) at representative cervical (C8), thoracic (T4) and lumbar levels (L4) from four samples. Arrowheads mark cell clusters in the LMC (black), PGC (gray) and HMC/MMC (white). Middle row: Overlay of VAChT^+^/Foxp1^+^ and ChAT^+^ cell positions, measured in one animal (#2). Bottom row: Overlay of VAChT^+^/Foxp1^+^ and ChAT^+^ cell positions, measured in two different animals (#3, #4). **F**) Relative distribution of VAChT^+^/Foxp1^+^ cells (red bars) and ChAT^+^ cells (blue bars) within motor columns of four animals (SAC: spinal accessory motor column, PMC: phrenic motor column). All VAChT^+^/Foxp1^+^ neurons express ChAT and represent 28.2% of ChAT^+^ LMC, 33.2% of ChAT^+^ PGC, 9.2% of ChAT^+^ MMC and 5.1% of ChAT^+^ HMC neurons.

#### Mapping different cell populations across samples

The precise mapping of cells from multiple animals using SpinalJ also provides for close comparison of the relative positions of different cell populations. To test and validate mapping of multiple populations and across samples, we plotted the positions of two different but overlapping sets of neurons: the cholinergic Foxp1 neurons (VAChT/Foxp1:tdTomato) and the entire cholinergic population of neurons (marked by anti-ChAT antibody). As expected, in four animals, we found that VAChT^+^/Foxp1^+^ cells lie within the distribution of the ChAT^+^ cell population (Fig.9E, top row). We compared the relative cell numbers within each motor column (Fig.9F) and calculated that VAChT^+^/Foxp1^+^ neurons account for subsets of ChAT^+^ LMC, PGC, MMC and HMC neurons. Notably, the relative distributions of VAChT^+^/Foxp1^+^ and ChAT^+^ cell populations labeled in the same animal versus in different animals matched closely (Fig.9E, middle and bottom rows), demonstrating the utility of SpinalJ for comparative analyses of relative spatial information in whole SC.

In summary, we provide a toolbox for the analysis of neurons and connections within the full r-c extent of mouse SC. We have developed SpineRacks for oriented embedding and efficient sectioning, SpinalJ for user-friendly image processing and analysis, and a 3D SC atlas that provides a standardized reference for analysis of spinal neurons and projections. We have validated the accuracy and reproducibility of SpinalJ mapping with reference to published manual studies, attesting to its usefulness for a variety of experiments. In addition, the availability of a common coordinate framework and 3D anatomical annotations for the first time permits comparative mapping of spinal neurons across samples and labs.

### Discussion

Given the technical limitations of traditional histological methods and more recent clearing approaches, SC analyses in mice have typically focused on sparse sampling through the entire SC or on subsets of spinal segments, resulting in incomplete characterization of the distributions and diversity of spinal neurons and their projections along the r-c axis as highlighted, for example, in Francius et al., 2013. Data generated by sparse sampling of whole SC is difficult to map accurately to defined r-c positions without the context of neighboring sections or vertebral landmarks (Harrison et al., 2013). In this study, we present tools to improve sectioning efficiency of whole SC and to facilitate section registration, atlas mapping and 3D analysis of SC sections. These tools fill the prevailing gap in methodology for SC analysis that limits detailed characterization of spinal cell types and circuitry. Moreover, mapping of section data to a common coordinate framework using SpinalJ permits, for the first time, SC wide analysis with high spatial precision and comparison of results from different samples and across research groups.

#### SpineRacks & Sectioning

We developed SpineRacks to overcome the massive effort of sectioning entire mouse SCs. While fully automated platforms exist to section and image tissue (Oh et al., 2014; Ragan et al., 2012), such systems are expensive, require specialist setup and maintenance, and are typically available only at dedicated research institutions. The methods presented here are inexpensive and can be readily applied in any lab with access to a cryostat. Using SpineRacks, >1000 25μm sections comprising spinal segments C1-L6 can be produced within 1h and collected on as few as 16 slides. Cutting thicker sections can further reduce sectioning and processing time at the expense of r-c resolution. To minimize damage and loss of information when cutting SC tissue for parallel embedding, careful placement of cuts orthogonal to the long axis of the SC is essential. Angled cuts produce incomplete transverse sections that cannot be registered and need to be discarded from analysis in SpinalJ. The Allen Institute developed special cutting tools using cast agar in the shape of the adult SC to support and orient unfixed tissue during cutting (Allen Institute for Brain Science, 2008). While these tools cannot be directly applied to samples that have been fixed *in situ*, due to the natural curvature of the SC, fabrication of an adapted mold and blade guide may further improve cutting precision. Alternatively, SC tissue can be straightened prior to fixation. In addition, to compensate for loss of data at cutting interfaces and to generate a complete dataset, cuts can be placed at slightly staggered intervals along the r-c axis of SC in different samples, allowing smooth analysis through breakpoints.

Using commercial PVA filament, SpineRacks are easy to produce on any fused deposition modeling (FDM) 3D printer on site, or through online 3D printing services. PVA is commonly used in biomedical products such as soft contact lenses, eye drops, drug tablets, embolization particles, and as artificial cartilage, due to the material’s high biocompatibility and low protein adsorption characteristics (Baker et al., 2012; Xu et al., 2019). It is thus unlikely that PVA would adhere to the tissue sections after washing slides in PBS and interfere with immunohistochemistry or fluorescence. Indeed, we have never observed adverse effects in a wide variety of experiments.

#### Imaging and registration

The precise arrangement of tissue segments embedded in the SpineRack makes it possible to automate image acquisition and segmentation of SC sections. We used a programmable slide scanning microscope to capture each block section on the slide automatically as a separate image file. Other devices may require manual selection of block section arrays or scanning of the entire slide. A major advantage of embedding tissue in SpineRacks is that r-c identity of individual tissue sections can be determined solely by their relative position within the block section array without the need to evaluate segment-specific morphological features. This enables fully automatic segmentation of tissue sections from block section images. Since our current segmentation approach is based on the relative spatial positions of tissue sections it is robust against distortion and deformation of sections. However, block sections that contain less than nine tissue sections cannot be processed automatically using this method and require simple manual placement of a segmentation mask.

For section registration, SpinalJ performs a rigid body registration of the reference channel (DAPI or NT). The quality of section registration critically relies on intact spinal sections. Images of damaged and incomplete sections, as well as out-of-focus images, cannot be used and must be replaced by neighboring sections. In the samples analyzed for this study, we had to replace up to 15% of sections distributed throughout the length of any given SC. Comparative analysis of multiple samples suggests that this intervention did not confound the real distribution of cells. Replacing larger numbers of sections may however substantially reduce axial resolution locally and introduce artifacts in cell distributions, especially when analyzing small, local cell clusters. Thus, good section quality is critical and tissue segment break points must be avoided in areas of particular interest. A tape transfer system like Leica’s CryoJane tape station may help to reduce deformation and damage to tissue sections.

#### Atlas and 3D annotations

SpinalJ provides the first 3D atlas and a common coordinate framework for mouse SC. The creation of this 3D atlas from 2D annotations of individual segments does not provide additional r-c resolution, but rather brings the existing annotations into r-c context and into a format that allows 3D registration and atlas mapping. It is important to note that this atlas and the Allen Spinal Cord Atlas annotations are based on sections of a single animal and can therefore not account for inter-animal variability in SC anatomy. In addition, while most of the larger annotated regions in the Allen Spinal Cord Atlas are symmetrical across the midline and drawn in both hemisegments, some of the smaller annotations (e.g. of MN clusters) appear only in one hemisegment, e.g. Ph9 in C3; Tz9, LS9, De9 in C4; Man9 in T2; etc.. These asymmetries in the annotations could have resulted from sections cut slightly obliquely or from incomplete staining and show that the annotations for a complete spinal segment cannot be derived reliably and with spatial precision from a single section.

To generate more stable annotations that are robust with regard to inherent anatomical variability and increase r-c resolution, additional annotation data from multiple samples are needed. However, to prepare, image and annotate SC sections representing the entire SC involves considerable effort and time. Also, in this scenario, assignment of segment boundaries is challenging and can be achieved precisely only with the help of additional physical landmarks e.g. nerve roots, as Nissl/AChE descriptors alone are inconclusive at some levels, especially in the thoracic SC (Harrison et al., 2013). Instead, we propose that 3D reconstruction of whole SCs and mapping of defined populations of neurons to a standard reference can be used to create and improve 3D annotations. Using Nissl, ChAT, and other cell type-specific markers, the boundaries of anatomical landmarks and specific cell clusters will emerge in their 3D shape and can be outlined directly to refine annotations and segment boundaries after mapping multiple samples to the common reference template in SpinalJ. Template mapping of registered sections relies solely on a continuous Nissl/NT reference and while, currently, the Nissl template used for mapping offers reduced detail to match the available annotations, improving template quality is much easier to achieve than acquiring new anatomical annotations in 2D sections. Providing our tools as an open resource, we hope progressively to improve the quality of the atlas with increasing numbers of datasets and markers mapped by individual labs and/or shared within the community. Moreover, additional 3D atlases, for example based on the annotations created for a P4 animal (Sengul et al., 2012), could be easily incorporated into SpinalJ.

#### Analysis in SpinalJ

Atlas mapping in SpinalJ is achieved by the registration of the DAPI or NT experimental volume to the 3D Nissl template. An affine 3D transformation has to be used to prevent deformation and warping of the experimental data during registration. With this, the width-to-length ratio of the experimental dataset is fixed and must match the template. However, loss of sections can shorten the experimental data along the r-c axis (the average shortening in the samples processed for this study was 39 sections = 975μm), resulting in mismapping. To compensate for this, SpinalJ matches the total r-c length of the experimental dataset to the length of the corresponding segment range in the template by interpolating additional sections. The current compensation approach assumes random/equal spacing of missing sections and does not account for locally concentrated losses, which can, however, be manually accommodated. With this approach, the accuracy of mapping NT signal to the GM annotation was determined to be 90% and 81% across the whole SC using NT or DAPI, respectively for section registration and template mapping (Fig.3). We observed these minor mismappings mainly at the transitions between the wide segments of the enlargements and the round segments of the thoracic SC, where slight r-c misalignment between experimental data and template would shift GM signals into WM annotations and vice versa. Additionally, using a single Nissl reference section as a mapping template for the entire length of each spinal segment does not account for slight anatomical variation within segments, thus introducing mapping imprecisions. Even more accurate r-c mapping could be achieved by introducing a continuous Nissl template and allowing for the specification of additional segment key frames at multiple r-c levels in future versions of SpinalJ.

Nonetheless, with the current version of the atlas, we have demonstrated that SpinalJ can reproduce the results of various manual mapping studies. In all validation datasets, we found good agreement of SpinalJ-mapped data and known distribution patterns within GM laminae and major WM tracts of isolated spinal regions. A few minor mapping errors were observed and typically resulted from signals shifted slightly across the original annotation outlines into neighboring annotations. For example, although AAV labeled CST axons mapped primarily to the dorsal corticospinal tract (dcs), there was minor mismapping to the annotation of the postsynaptic dorsal column pathway (psdc), located immediately dorsal to the dcs (Fig.6D,G). For smaller annotation regions, like the individual MN clusters of lamina IX that cover only a few neurons, the mapping errors appeared larger. Mapping VAChT^+^/Foxp1^+^ neurons, we identified significant signal outside lamina IX, in neighboring areas lamina VII and VIII (Fig.S5E). Similarly, in samples stained for IB4^+^ fibers, we found some mismapping to neighboring laminae outside lamina II (Fig.5F). Improving the quality and resolution of the atlas annotations will address these minor issues of mapping imprecision. Indeed, irrespective of annotation boundaries, there was a high mapping precision of the positions of cell populations across different animals. Measuring centroid distances between animals, we determined an average mapping offset of only 13μm for ChAT^+^ neurons and 31μm for VAChT^+^/Foxp1^+^ neurons, i.e in the range of a single MN cell body diameter. Mapping appears slightly more precise for ChAT^+^ neurons because centroid positioning is more robust with higher cell numbers.

Thus, with minimal inter-animal mapping offsets, SpinalJ is well suited to analyze the relative positions of known and unknown cell types and to map cells and circuits across samples, which has not been possible previously. This is illustrated by the analysis of VAChT^+^/Foxp1^+^ neurons that revealed an unexpected population of cells in the ventral thoracic SC (Fig.8). Based on their position, the labeled neurons could belong to the pool of MMC or HMC MNs, although this lineage is defined by the embryonic expression of Lhx3 and the suppression of Foxp1 expression, likely through direct co-regulation of both factors (Morikawa et al., 2009). Within the embryonic thoracic SC, Foxp1 was detected only in Isl1^+^/pSmad^+^ PGC neurons (Dasen et al., 2008; Morikawa et al., 2009). It is, therefore, unlikely that the labeled cells belong to MMC or HMC MNs, unless these cells start expressing Foxp1 at later stages, something that has not been examined to date. Outside the population of MNs, Foxp1 expression has been observed in Pax2^+^/En1^+^ V1 interneurons during mid- to late embryonic stages (Francius et al., 2013; Morikawa et al., 2009). These cells are positioned close to ventral MNs and generally fit the observed distribution. However, V1s have been described as inhibitory GABAergic or glycinergic neurons, excluding ChAT immunoreactivity, unless at least some V1s co-transmit acetylcholine and GABA, as has been observed in other neurons (Lamotte d’Incamps et al., 2017; Vaaga et al., 2014). Within the group of ventral interneurons, only Pix2^+^ V0c neurons have been identified as cholinergic (Zagoraiou et al., 2009; Ziskind-Conhaim and Hochman, 2017), but Foxp1 does not co-localize with V0 markers embryonically (Morikawa et al., 2009). Further analyses and co-staining with markers for ventral interneuron classes are needed to identify the labeled cell population identified by our study.

Standardized 3D anatomical atlases, such as the Allen Brain Atlas, generated from iterative averaging of over 1500 different mouse brain samples (Wang et al., 2020), have proven an indispensable resource in brain research. Atlases provide a high resolution framework for comparative analyses and have enabled massive collaborative projects like the BRAIN initiative (Ecker et al., 2017). Various computational tools have been developed for the analysis and integration of individual brain datasets (Bakker et al., 2015; Botta et al., 2020; Chon et al., 2019; Eastwood et al., 2019; Friedmann et al., 2020; Oh et al., 2014; Puchades et al., 2019; Shiffman et al., 2018; Tappan et al., 2019; Tyson et al., 2020; Wang et al., 2021), improving both speed and quality of experiments. Together, these resources have enabled brain-wide mapping studies of cell types and neuronal connectivity and dramatically accelerated scientific discovery in this field of research. In contrast, SpinalJ is the first toolbox for the 3D analysis of SC data in the context of anatomical annotations. We have demonstrated that, even in its prototype form, SpinalJ provides a powerful platform for high-throughput analysis of the relative positions of populations of genetically, virally or immunohistochemically labeled neurons, their projection patterns and axonal tracts. We aim to establish SpinalJ as a continuously improving resource for the field of SC research.

## Acknowledgements

We thank Samaher Fageiry for technical advice on SC tissue preparation and histology, Marcela Carmona and Anna Kim for help with cortical virus injections, Crystal K. Colón Ortiz for providing mouse eye samples, as well as Amy Norovich and Young Mi Kwon for providing betta fish brains. We thank Susan B. Morton and the Jessell lab for gifts of custom antibodies. We are grateful to Carl E. Schonoover and Sarah Ohashi for help with 3D printing and thank Isobel Jessell for the term ‘SpineRack’. We thank the Mason-Dodd and Bendesky labs for discussion and Christoph Gebhardt for critical comments on the manuscript. Imaging was performed with support from the Zuckerman Institute’s Cellular Imaging platform. This work was funded by NIH grants 1R21NS120665-01, 5U19NS104649 and DFG grant FI 2367/1-1.

## Author contributions

The study was conceptualized and directed by F.F., J.D. and C.M. F.F. developed SpineRacks and performed histology studies, imaging and data interpretation. L.A.H. and F.F. developed SpinalJ software. D.N. provided VAChT/Foxp1 mice. F.F. and J.D. wrote the manuscript with input of all authors.

## Declaration of interests

Authors declare no competing interests.

## STAR methods

### Resource availability

#### Lead contact

Further information and requests for resources and reagents should be directed to and will be fulfilled by the lead contact, Jane Dodd.

#### Materials availability

This study did not generate new unique reagents.

#### Data and code availability

The datasets and code generated during this study are available at Github (https://github.com/felixfiederling/SpinalJ).

### Experimental models and subject details

#### Animals

All experimental protocols were approved by the Columbia University Institutional Animal Care and Use Committee. All experimental animals were adult (>3 month old) male and female mice housed on a 12h light/dark cycle with *ad libitum* access to food and water. Unless stated otherwise, we used C57BL/6J animals for tool development and validation. To label cholinergic, Foxp1 expressing neurons, we used *FoxP1::FonCre^FonCre/+^*; *VAChT::FlpO^FlpO/+^; Ai9^A19/+^* mice. In *FoxP1::FonCre^FonCre/+^* mice, Cre recombinase is incorporated at the 3’ end of *FoxP1* using a P2A linker to avoid disruption of *FoxP1* expression. The Cre open-reading-frame is interrupted by a ‘stop cassette’ flanked by F3 FRT sequences, and is restored with Flp expression. Both *FoxP1::FonCre^FonCre/+^* and *VAChT::FlpO^FlpO/+^* mice were generated by homologous recombination using mouse ES cells. The details of these two mouse lines will be presented elsewhere (Ng et al., in preparation).

### Method details

#### 3D printing

SpineRack embedding scaffolds were designed using Autodesk TinkerCAD (www.tinkercad.com; Autodesk, Inc) and printed from Ultimaker Polyvinyl alcohol (PVA) filament (Ultimaker B.V.) on a dual extruder Ultimaker 3 printer. Ultimaker Cura software was used for slicing and printer setup (material: natural PVA; print core: BB 0.4; layer height: 0.15mm; print temp: 220°C, bed: 60°C; infill: 20%; build plate adhesion: brim 3mm).

For all results shown, we used SpineRacks with outer dimensions: 11mm x 11mm x 4.0mm, well size: 3.0mm x 3.0mm x 4.0mm, wall thickness: 0.5mm. Other dimensions are easily achieved. Print files are available for download (see Key Resources Table).

PVA is hygroscopic and to prevent absorbance of moisture from room air filament and printed racks should be stored in the dark in an air tight container along with a desiccant. Under these conditions, we have found that SpineRacks can be stored for at least one year without qualitative changes, swelling or shrinking.

#### Viral labeling of corticospinal neurons

Cortical virus injections were performed under sterile conditions and isoflurane anesthesia (1–3%, plus oxygen at 1-1.5L/min) on a stereotactic frame (David Kopf Instruments, Model 900SD). Throughout surgery, mouse body temperature was maintained at 37°C using an animal temperature controller (FHC, Model 40-90-8D) and, afterward, mice were allowed to recover from the anesthesia in their homecage on a heating pad. Before surgery, animals were subcutaneously injected with Buprenorphine SR (0.5-1mg/kg). The mouse head was shaved, cleaned with 70% alcohol and iodine, an intradermic injection of bupivacaine was administered and the skull was exposed to permit alignment of the head and drilling of the hole for the injection site. 500nl of AAV2.1-CAG-TdTomato (titer: 5.3×10^12^ vg/ml; UNC, lot AV6325C) was injected across five injection sites into the left hemisphere of the motor and sensory cortex (central coordinate: AP -.25mm, ML 1.5mm, and DV between .45-.85mm with four additional injections spaced ~500μm apart, forming a square around the center) using a Nanojet III Injector (Drummond Scientific, USA) at a pulse rate of 1nl/s, injecting 20-25nl every 100-200μm. The injection pipette was left in place for 10min post-injection before it was slowly removed (rate 200μm/s). After injections, the small whole made during the craniotomy was filled with kwik-sil silicon adhesive (World Precision Instruments, USA) and the skin was closed using sutures. After 4 weeks, mice were euthanized and perfused as described below. Brains were embedded in 2% agarose and sectioned on a vibratome at 100μm. To determine the position and extent of injection sites, images of brain sections were registered and mapped to the Allen Mouse Brain Atlas using BrainJ, as described in (Botta et al., 2020).

#### Tissue preparation

Mice were perfused with 4% paraformaldehyde (PFA) in PBS and the spinal column was post-fixed in 4% PFA overnight at 4°C after exposing the SC through ventral laminectomy. The SC was then isolated, washed 3x in cold PBS and cryo-protected in 30% sucrose solution at 4°C until the tissue had sunk. To optimize sectioning efficiency of the whole SC, nine sequential tissue pieces were mounted in one block. For this, the SC was first trimmed caudally, removing segments caudal to S1 and the *cauda equina*. We then split the remaining cord, spanning all cervical, thoracic and lumbar segments, into three equal sized tissue pieces using iridectomy scissors. Each of these pieces was then split again into three equal sized pieces, resulting in a total of nine, 3-4mm long tissue pieces (Fig.1A-D; for detailed instructions, see supplementary user guide).

#### Embedding

For embedding, a truncated 12mm plastic mold (Peel-A-Way T12; Polysciences, Inc.) was filled with Tissue-Tek OCT Compound (Sakura Finetek USA, Inc.). A SpineRack was then sunk into the OCT and pushed to the bottom of the mold using blunt forceps. Air bubbles trapped in the structure were removed using the sharp points of forceps. Spinal tissue pieces were gently placed into each well, until the rostral cut face touched the bottom of the mold. Pieces 1 to 3 were embedded (left to right) in the top row, pieces 4 to 6 in the middle row and pieces 7 to 9 in the bottom row of the rack (Fig.1E-I). Other tissues (fish brains, mouse eyes) were embedded similarly. To facilitate easy cryo-sectioning through the block containing SpineRack and tissue pieces, the filled mold was left at room temperature for 20-25min before freezing. This allowed the SpineRacks partially to dissolve and soften in the OCT and to achieve consistent texture across the block. Filled molds were then frozen on dry ice in a slush of absolute ethanol and crushed dry ice and resultant blocks stored at −80°C until use.

#### Sectioning and Immunohistochemistry

Blocks were sectioned on a Leica CM3050S cryostat (Leica Biosystems) at 25μm. Sections were collected on Fisherbrand Superfrost Plus slides (Fisher Scientific). Eight consecutive sections were collected in two rows on each slide (Fig.1J-K). Slides were washed in Wheaton staining dishes filled with PBS, for 5 min on an orbital shaker, to dissolve OCT and SpineRack material. Sections were incubated with primary antibodies in PBS containing 0.1% TritonX-100 at 4°C overnight, then washed in PBS and incubated with secondary antibodies, DAPI and/or Neurotrace in PBS for 1-2h at room temperature. tdTomato signal was amplified using anti RFP or anti dsRed antibodies. See Key Resources Table for material details.

#### Imaging

Slides were imaged using a motorized Nikon AZ100 Multizoom microscope (Nikon Instruments Inc.) equipped with an automated slide feeder (Prior Scientific Inc.) and Andor Zyla sCMOS camera (Oxford Instruments). Images were acquired using a Nikon 4x 0.4 NA AZ Plan Apo objective with an additional 2.1x magnification, resulting in an image pixel size of 1μm/pixel. Each block section (3×3 array of nine tissue sections) was scanned and saved as one image file (.nd2 format, containing stage coordinate metadata) using NIS-Elements JOBS software (configuration file available for download at; see Key Resources Table). To do this, the software was programmed to scan the entire slide at low resolution in a single channel (DAPI or NT). A manually determined threshold was then applied automatically to isolate tissue sections from background and a dilation factor was used to add pixels to the object boundaries and merge all tissue sections of a block section into a single object. The identified block section objects were then scanned in all channels. This approach allowed us to keep the positional information of tissue sections within a block section, which is essential to identify individual tissue sections, while keeping the file size of images in a manageable range (1-2GB per block section image). We imaged a single optical plane of each 25μm section, assuming that there would be minimal overlap of cell somata in the z direction with spinal cell types ranging from 7-45μm in diameter (Sengul et al., 2012).

#### Image processing and analysis

Images of SC tissue sections were processed and reconstructed in Fiji/ImageJ (Schindelin et al., 2012) using SpinalJ, a plugin developed in this study combining software tools to facilitate image registration, atlas mapping and 3D analysis of SC sections. SpinalJ and a detailed, step-by-step user guide are freely available for download (see Key Resources Table).

For easy visualization of data, SpinalJ creates plots of absolute and relative signal intensities, cell densities and projection densities as heatmap montages for each spinal segment. In addition, we used the 3D viewer plugin for Fiji (Schmid B. et al., 2010) to render 3D views of reconstructed datasets. 2D/3D cell position plots were created in MATLAB (Mathworks) using the *‘scatter’* and *‘scatter3’* functions, respectively. Similarly, heatmap chart matrix plots were created in MATLAB using the *‘heatmap’* function.

All image processing and analysis was performed on a workstation running Windows 10 Enterprise 64 bit, equipped with a 16 core Intel i9 7960x 2.8GHz CPU, 128 GB DDR4 memory, a 1TB Samsung 860 SSD and a 12GB Nvidia Titan X video card. SpinalJ processing generates a significant amount of data per dataset (2-3 fold original data) to allow for validation of results and reprocessing when required, but these intermediate data can be deleted following successful processing.

### Quantification and statistical analysis

All values are shown as mean ± standard deviation of the mean (SEM). Statistical details of experiments are provided in figure legends.

### Key Resource Table

| REAGENT or RESOURCE | SOURCE | IDENTIFIER |
| --- | --- | --- |
| **Antibodies** |  |  |
| Rabbit polyclonal anti-RFP | Rockland Immunochemicals | Cat#600-401-379 |
| Guinea pig polyclonal anti-dsRed | Gift of Jessell lab | N/A |
| Goat polyclonal anti-ChAT | Millipore Sigma | Cat#AB144P |
| **Bacterial and virus strains** |  |  |
| AAV2.1-CAG-tdTomato | UNC vector core | Lot#AV6325C |
| **Chemicals, peptides, and recombinant proteins** |  |  |
| Isolectin B4, FITC-conjugated | Sigma-Aldrich | Cat#L2895 |
| Neurotrace 500/525 | Thermo Fisher Scientific | Cat#N21480 |
| Neurotrace 640/660 | Thermo Fisher Scientific | Cat#N21483 |
| DAPI | Thermo Fisher Scientific | Cat#D1306 |
| **Deposited data** |  |  |
| SpineRack and SpinalJ user guide | This study | https://github.com/felixfiederling/SpinalJ/blob/main/SpineRack%20and%20SpinalJ%20User%20Guide.pdf |
| 3D SC reference atlas p56 | This study | https://data.mendeley.com/datasets/4rrggzv5d5/1 |
| Raw data and analysis spreadsheets | This study | https://github.com/felixfiederling/SpinalJ/tree/main/data |
| **Software and algorithms** |  |  |
| Fiji (ImageJ) | NIH | https://fiji.sc/ |
| SpinalJ | This study | https://github.com/felixfiederling/SpinalJ |
| Ilastik 1.3.3 | Berg et al., 2019 | https://www.ilastik.org/ |
| Matlab | Mathworks | https://www.mathworks.com/products/matlab.html |
| NIS-Elements JOBS | Nikon Instruments | https://www.microscope.healthcare.nikon.com/en_EU/products/software/nis-elements/nis-elements-jobs |
| JOBS settings for imaging | This study | https://github.com/felixfiederling/SpinalJ |
| **Other** |  |  |
| PVA filament | Ultimaker | https://ultimaker.com/materials/pva |
| SpineRack .stl print file | This study | https://github.com/felixfiederling/SpinalJ/blob/main/SpineRack.stl |

## Notes

https://github.com/felixfiederling/SpinalJ

https://data.mendeley.com/datasets/4rrggzv5d5/1

### Competing Interest Statement

The authors have declared no competing interest.

