## Supplemental Figures for "Tools for efficient analysis of neurons in a 3D reference atlas of whole mouse spinal cord"

Figure S1

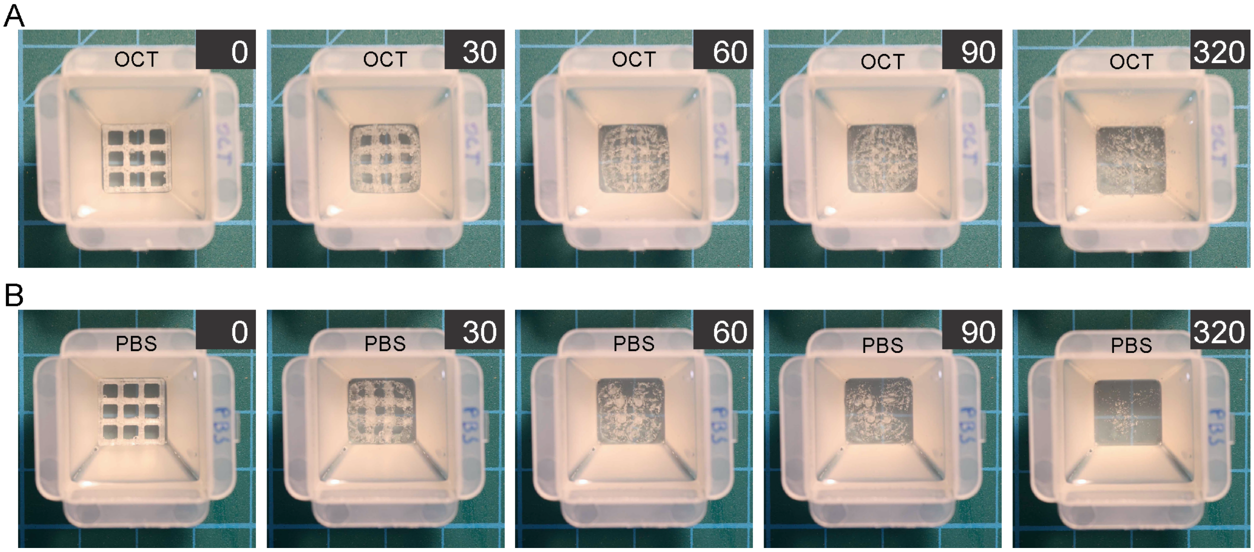


Figure S1: SpineRack solubility
PVA SpineRacks dissolving in OCT (**A**) or PBS (**B**) at room temperature. Time in minutes.

Figure S2

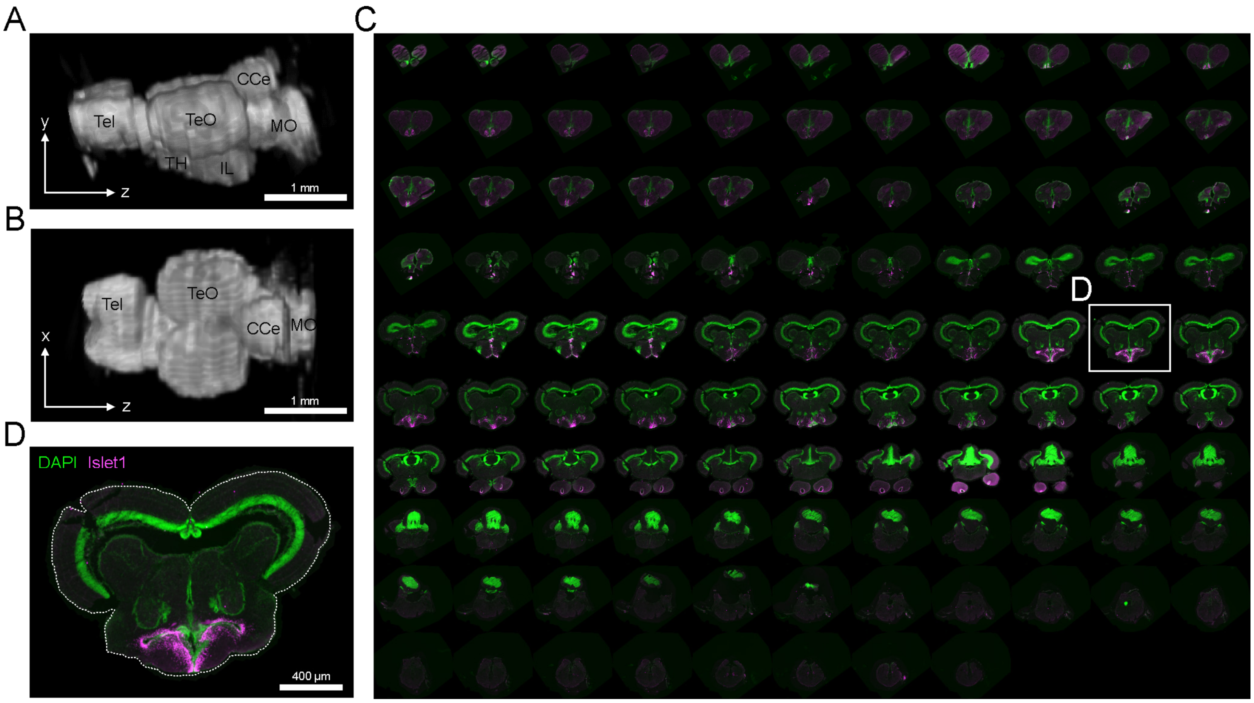


<u>Figure S2: SpineRacks for oriented sectioning of *Betta splendens* brains</u>
Four brains of adult *Betta splendens* were embedded using a SpineRack for synchronous sectioning. Reconstructed sections from one brain shown in lateral (**A**) and dorsal (**B**) views after registration in SpinalJ. Tel: Telencephalon; TeO: Tectum opticum; CCe: Corpus cerebelli; MO: Medulla oblongata; TH: Tuberal hypothalamus; IL: Inferior lobe of hypothalamus. **C**) Montage of the ordered stack of sections counterstained for Islet1 (magenta) and DAPI (green). **D**) Magnified view of section highlighted in C.

Figure S3

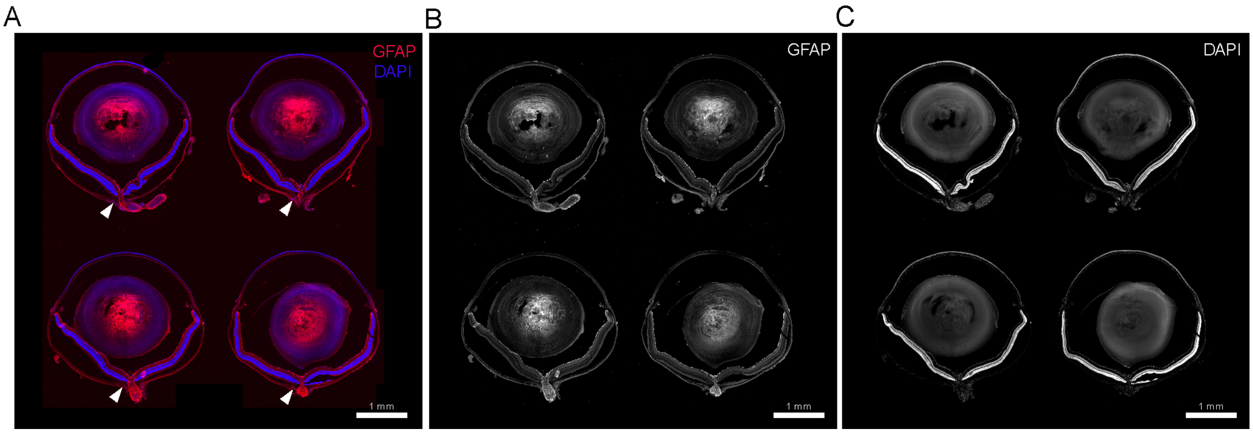


Figure S3: PVA racks for oriented sectioning of mouse eyes

**A**) Block section at the level of the optic nerve (arrowheads) of four adult mouse eyes (two animals) that were oriented and embedded in a modified SpineRack offering four, 4.7 x 4.7mm wide wells. Section is stained for GFAP (red) and DAPI (blue). **B**) GFAP channel of image in A. **C**) DAPI channel of image in A.

Figure S4

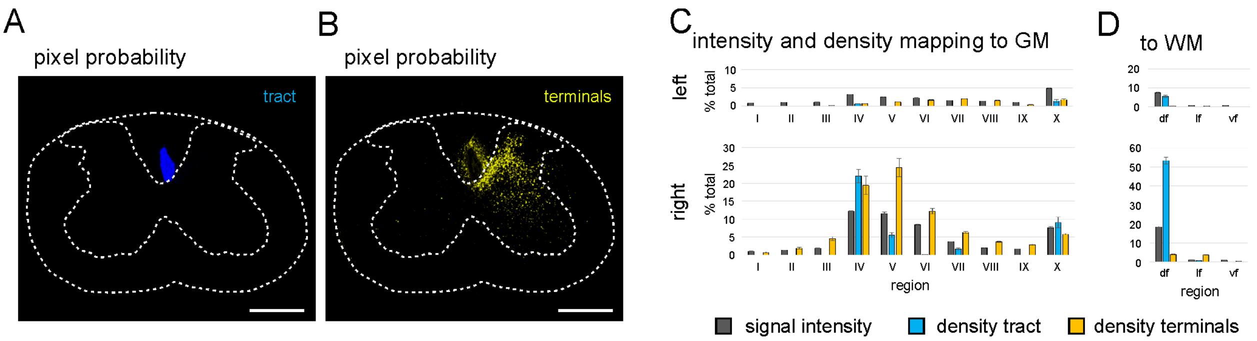


Figure S4: Segmentation of tract and terminal/branch signals

**A**, **B**) Pixel probabilities for classifiers 'tract' (A, blue) and 'terminals/branches' (B, yellow) after training in Ilastik. Scale: 500µm. **C**, **D**) Relative distribution of signal intensities (gray), and densities of segmented tract (blue) and terminal/branch signals (yellow) within atlas regions of the GM (laminae I-X) and WM (df, lf, vf) of the left, ipsilateral (top row), and right, contralateral (bottom row), hemisegments of C4-C7. Error bars indicate standard deviation of values from all segments within the analyzed range.

Figure S5

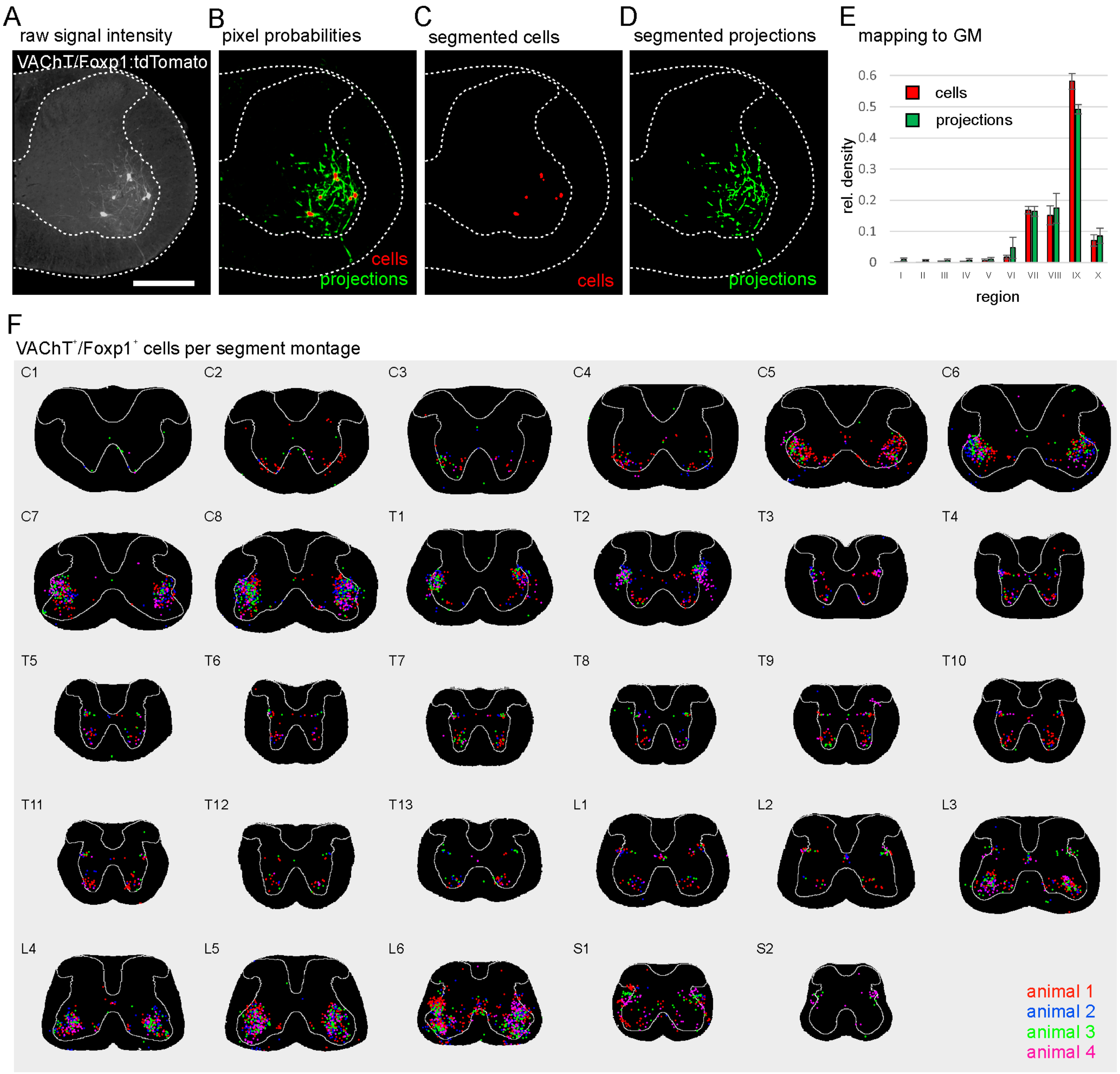


Figure S5: Detection and mapping VAChT$^+$/Foxp1$^+$ cells and projections

**A**) tdTomato expression in a SC section of a VAChT/Foxp1 animal. Scale bar: 500µm. **B**) Pixel probabilities for classifiers 'cells' (red), 'projections' (green) after training in Ilastik. **C**, **D**) Cells and projections detected after image segmentation using pixel probabilities. **E**) Relative distribution of mean VAChT$^+$/Foxp1$^+$ cell (red) and projection densities (green) of four animals within atlas regions of the GM (laminae I-X). Error bars indicate standard deviation between values from both hemisegments. **F**) 2D distribution of VAChT$^+$/Foxp1$^+$ cells from different samples within each spinal segment.
